# Induced microfold cells are required for the function of pulmonary tertiary lymphoid organs that promote mucosal immunity

**DOI:** 10.64898/2026.10.28.762535

**Authors:** Manalee Vishnu Surve, Dawei Sun, Vanshika Panchal, Brian Lin, Po Kwok Tse, Sunghyun Kim, Qiaozhen Liu, Tivadar Török, Jackson A. Weir, Anthony Xu, John Vo Huynh, Victoria I. Cottrell, Hannah Willit, Giovanni Marrero, Vipin Kumar, Jiawei Sun, Steven Zwick, Viral S Shah, Jason W. Griffith, Fei Chen, Jayaraj Rajagopal

## Abstract

Tertiary lymphoid organs (TLOs) arise in response to infection and inflammation. We show that influenza-induced lung TLOs are comprised of inducible M cell (iM cell)-inducible bronchus-associated lymphoid tissue (iBALT) functional composites. iM cells are necessary for the appropriate architecture of adjacent immune follicles, which contain clonally expanded and sequence-diversified B cells. iM cell ablation results in fewer antigen-specific germinal center B cells, T follicular helper cells, and T helper memory cells, and a decrease in secretory IgA and IgG production accompanied by a loss of viral control. We find that the iM cells arise from epithelial progenitors that respond to a RANKL signal from infiltrating T cells, consistent with a model in which the patchy inflammation typical of pneumonia results in the unpredictable and varying distribution of coupled iM cells and adjacent iBALT. We propose that iM cells could be harnessed to promote infection control and enhance mucosal vaccination.

## Introduction

Tertiary lymphoid organs (TLOs) are organized lymphoid structures that arise in the setting of infections and autoimmunity, and their presence in cancers has recently engendered great interest. TLO architecture closely resembles that of secondary lymphoid organs (SLOs), but the detailed biology of TLO induction and function has only recently come under scrutiny.^1–3^ In the adult lung, TLOs arise in the form of inducible bronchus-associated lymphoid tissue (iBALT) in response to chronic inflammation and infection. Gut-associated lymphoid tissues (GALT), classified as SLOs, were described as early as 1677 given that Peyer’s patch follicles are an ever-present and easily recognized histologic feature of the steady-state intestine.^4^ In contrast, as it is not a stereotyped feature of the steady state adult lung, BALT had escaped attention until much later.^5^ Nonetheless, iBALT recapitulates the key architectural features of SLOs, including segregated B- and T-cell domains, germinal centers, follicular dendritic cell networks, and specialized vasculature known as high endothelial venules that together support local antigen presentation, germinal center reactions, the generation of memory, and protective anti-viral mucosal immunity.^6–9^

The principles of mucosal immunity have largely been defined through the study of GALT. GALT comprises lymphoid follicles that are intimately associated with a specialized intestinal follicle-associated epithelium (FAE). Furthermore, the FAE contains microfold (M) cells that transcytose luminal antigens and microbes through the gut barrier to the underlying immune cells of the adjacent follicle, ultimately engendering a robust antigen-specific mucosal immune response.^10^ Notably, the adult lung was not known to harbor resident M cells until Kimura and colleagues recently identified the presence of rare lung M cells.^11^ In the setting of infection and inflammation, greater numbers of inducible M cells (iM cells) become physically approximated to iBALT.^11,12^ Most recently, we reported the first airway M cell transcriptomes and demonstrated that airway M cells, like their gut counterparts, possess the ability for antigen and microorganism uptake.^11,13^ Furthermore, we have shown that airway M cells differentiate in response to inflammatory cytokines *in vitro.*^13^

Herein, using *in vivo* lineage tracing and genetic loss-of-function studies, we establish that RANK receptor-expressing secretory epithelial progenitors are the cellular source of influenza-induced lung iM cells. In embryonically patterned GALT, RANKL-expressing mesenchymal cells located between immature follicles and FAE promote the local differentiation of M cells in the overlying gut epithelium, thereby functionally coupling M cells with intestinal Peyer’s patches.^14,15^ Consistent with the varying localization of iBALT along the bronchial tree, we find no evidence for pre-patterned RANKL expressing mesenchymal cells in the lung, despite demonstrating the necessity of RANKL for lung iM cell differentiation. To identify the cellular source of RANKL, we employed our recently described Slide-tags single-nucleus spatial transcriptomics platform. Using spatial ligand–receptor analysis, we established the presence of multiple reciprocal signaling networks between iM cells and the immune cells of the adjacent follicles. We now find that infiltrating Th17, Tfh, and Treg cells are the key sources of RANKL in the lung, consistent with the patchy distribution of inflammation, characteristic of lung infection.^16,17^ In order to describe how iM cell–iBALT units foster mucosal immunity, we developed a methodology for coupling spatial BCR profiling with Slide-Tags. This allowed us to demonstrate active B cell clonal expansion and diversification within iM cell-associated follicles. Furthermore, the genetic ablation of M cells disrupted iBALT development and organization. Functionally, M cell ablation compromised iBALT-mediated antigen-specific germinal center B-cell responses, T follicular helper cell responses, and tissue-resident memory T cell responses. Most importantly, M cell loss resulted in compromised local IgA and IgG recall responses and viral control following heterotypic influenza challenge.

## Results

### Spatial transcriptomics reveals the cellular composition and architecture of lung M cell-associated tertiary lymphoid organs

To define the cellular neighborhood surrounding lung M cells, we employed Slide-tags,^18^ a state-of-the-art single-nucleus spatial transcriptomic methodology, and coupled it with our recently published M cell-specific *Sox8-CreER* driver mouse to allow M cell-specific labeling.^19^ We validated the labeling efficiency and specificity of this driver using *Sox8-CreER::LSL-tdTomato* reporter mice that had been treated with RANKL to induce M cell differentiation (Figure S1A). Tamoxifen administration resulted in the faithful epithelial labeling of >70% of mature CCL9⁺GP2⁺ double-positive M cells with >96% specificity (Figures S1B-S1D).^13,20^ For Slide-tags, we sectioned fresh-frozen lung tissues collected 2– 4 weeks after influenza A virus/PR8 infection of *Sox8-CreER::LSL-tdTomato* reporter mice. This enabled the detection of tdTomato-labeled M cells and their associated lymphoid aggregates. The adjacent sections were then spatially profiled using Slide-tags (Figure 1A).

**Figure 1.**
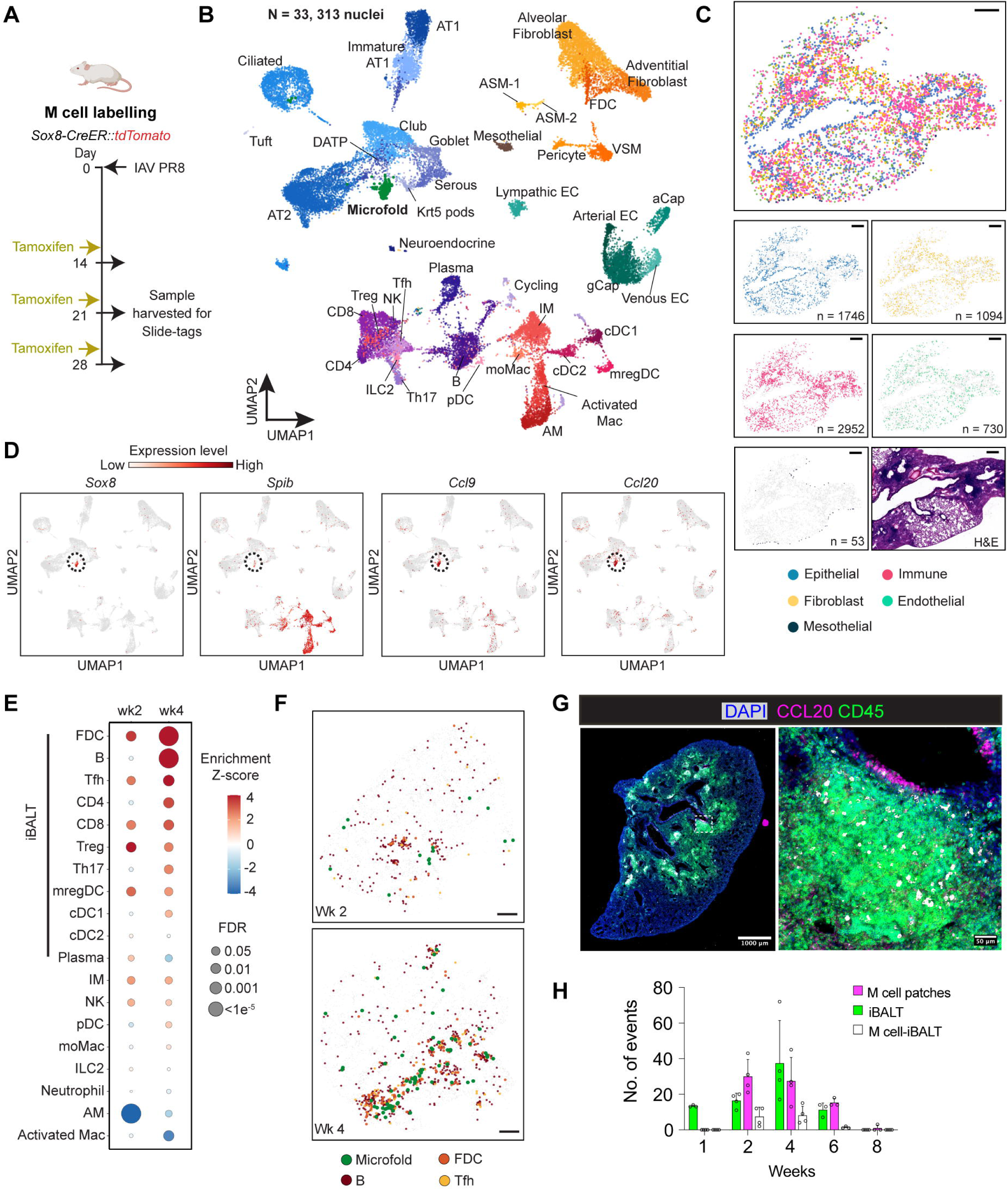
Spatial organization and cellular composition of M cell-associated iBALT in the influenza-infected lung. **A.** Schematic of the experimental workflow for Slide-tags profiling of influenza A-infected mouse lungs. **B.** UMAP representation of the time-resolved spatial transcriptomic dataset. **C.** Representative cell type-specific spatial maps and an adjacent haematoxylin and eosin (H&E)-stained section. Cells are colored according to cell type annotation. Data are shown for a 3-week post influenza A-infection sample. Scale bars, 500 μm. **D.** Representative expression patterns of M cell marker genes visualized in UMAP space. **E.** Dot plots of spatial co-enrichment of M cells with neighboring immune cell types at 2 week and 4 week post-influenza A infection. **F.** Spatial distribution of M cells alongside B, FDCs and Tfh cells in the representative lung tissue sections at 2 weeks (top panel) or 4 weeks (bottom panel) post-influenza A infection. Scale bars, 500 μm. **G.** Representative immunofluorescence image of lung section from wild-type mouse following influenza A infection. The region outlined by the dotted square in the whole-lung section is shown at higher magnification in the right inset. Nuclei: DAPI (blue). M cells: CCL20 (magenta). Immune follicles are identified by CD45⁺ immune cells (green) and GL7⁺ germinal center cells (gray). Scale bars, 1,000 µm (whole-lung section) and 50 µm (inset). **H.** Quantification of CCL20-positive M cell patches (>5 CCL20 positive cells), CD45 and GL7 positive iBALT structures, and M cell-associated iBALT in mouse lung sections at the indicated time points following influenza A/PR8 infection. Quantifications were averaged across two representative sections per mouse. (n = 3–4 mice per time point). Error bars indicate standard deviation (SD).

The resulting dataset consists of 33,313 high-quality and spatially resolved single nuclei and captures diverse epithelial, immune, fibroblast, endothelial, and mesothelial cell types (Figures 1B-1C and S2A-S2B). Cell identities were assigned based on established lineage- and cell state-specific transcriptional programs.^21,22^ M cells formed a distinct transcriptional cluster characterized by the enrichment of the canonical M cell markers including the lineage-specifying transcription factors *Sox8* and *Spib* and M cell-associated chemokines *Ccl9* and *Ccl20* (Figure 1D).^13,20^

We next examined how the M cell microenvironment evolves over the course of influenza infection. To characterize the cellular niches within the lung, we performed neighborhood enrichment analysis. Unsupervised clustering identified three spatially distinct cellular niches corresponding to the alveoli (enriched for AT1 cells, AT2 cells, and alveolar macrophages), airways (enriched for secretory and ciliated cells), and iBALT (enriched for M cells and diverse populations of T, B, and dendritic cells) (Figure S2C). At week 2 following infection, M cells were preferentially associated with follicular dendritic cells (FDCs), T follicular helper (Tfh) cells, CD8 T cells, and T regulatory cells (Tregs), consistent with the early assembly of induced lymphoid aggregates. By week 4, characteristic mature iBALT structures containing B cells, CD4 helper T cells, Th17, and cDC1 cells emerged.^7,9,23,24^ Additionally, FDCs were further enriched, as would be expected (Figures 1E-F and S2D-E).

To confirm these findings, we performed parallel immunofluorescence staining. Consistent with our spatial analysis, CD45⁺GL7⁺ germinal center (GC)-containing immune follicles were most abundant at 4 weeks post-infection (Figure 1H). This peak coincided with maximal M cell abundance. Furthermore, M cells occurred in patches that were closely associated with GC-containing immune follicles (Figure 1G), consistent with the prior report that established a physical correlation of M cells with iBALT.^12^ Thereafter, the abundance of M cell patches, GC-containing follicles, and M cell-associated follicles declined in parallel as M cell–iBALT units resolved (Figure 1H). Collectively, these analyses establish the first high-resolution, spatiotemporally resolved map of post-influenza M cell-associated iBALT.

### T cells are the source of RANKL necessary for M cell differentiation

RANKL–RANK signaling is essential for M cell differentiation in various mucosal tissues.^25–27^ Furthermore, we and others have previously shown that RANKL specifically induces functional airway M cells in primary epithelial cultures and promotes M cell differentiation in the lungs.^12,13^ To demonstrate whether RANK signaling is necessary for influenza-induced lung M cell differentiation and, furthermore, identify the cell populations producing RANKL in infected lungs, we examined the expression of *Tnfsf11* (RANKL) and its receptor *Tnfrsf11a* (RANK) across cell types in our spatial transcriptomic dataset.

In the small intestine, RANKL is constitutively expressed by specialized mesenchymal cells and is necessary and sufficient for the stereotyped patterning of M cells within the follicle-associated epithelium (FAE).^14,15^ In contrast to the gut, the mature lung does not possess lymphoid follicles. Instead, post-infectious iBALT forms in varying sites along the bronchial tree.^8^ Indeed, the patterning mechanisms governing TLO localization are poorly understood. We find that in the context of influenza infection, RANKL is expressed exclusively in lung T cell populations, including Tfh, Th17, and Tregs. On the contrary, RANK itself was broadly expressed in various epithelial, immune, fibroblast, and endothelial cell populations but most highly in M cells and presumptive mregDCs (Figure 2A). Spatial mapping further revealed that RANKL-expressing T cells preferentially localized in close proximity to RANK-expressing M cells, nominating Tfh, Th17, and Tregs as candidate cellular regulators of lung M cell differentiation (Figure 2B). We next assessed whether RANKL signaling is required for lung M cell differentiation by administrating RANKL-blocking antibody following influenza infection (Figure 2C). Anti-RANKL treatment abrogated M cell differentiation (Figure 2D), demonstrating that RANKL signaling is necessary for lung M cell differentiation during influenza infection. Together, these findings identify T cells as potential M cell inducers (MCi) in the lung, analogous to the mesenchymal MCi that support M-cell differentiation in Peyer’s patches.^14^

**Figure 2.**
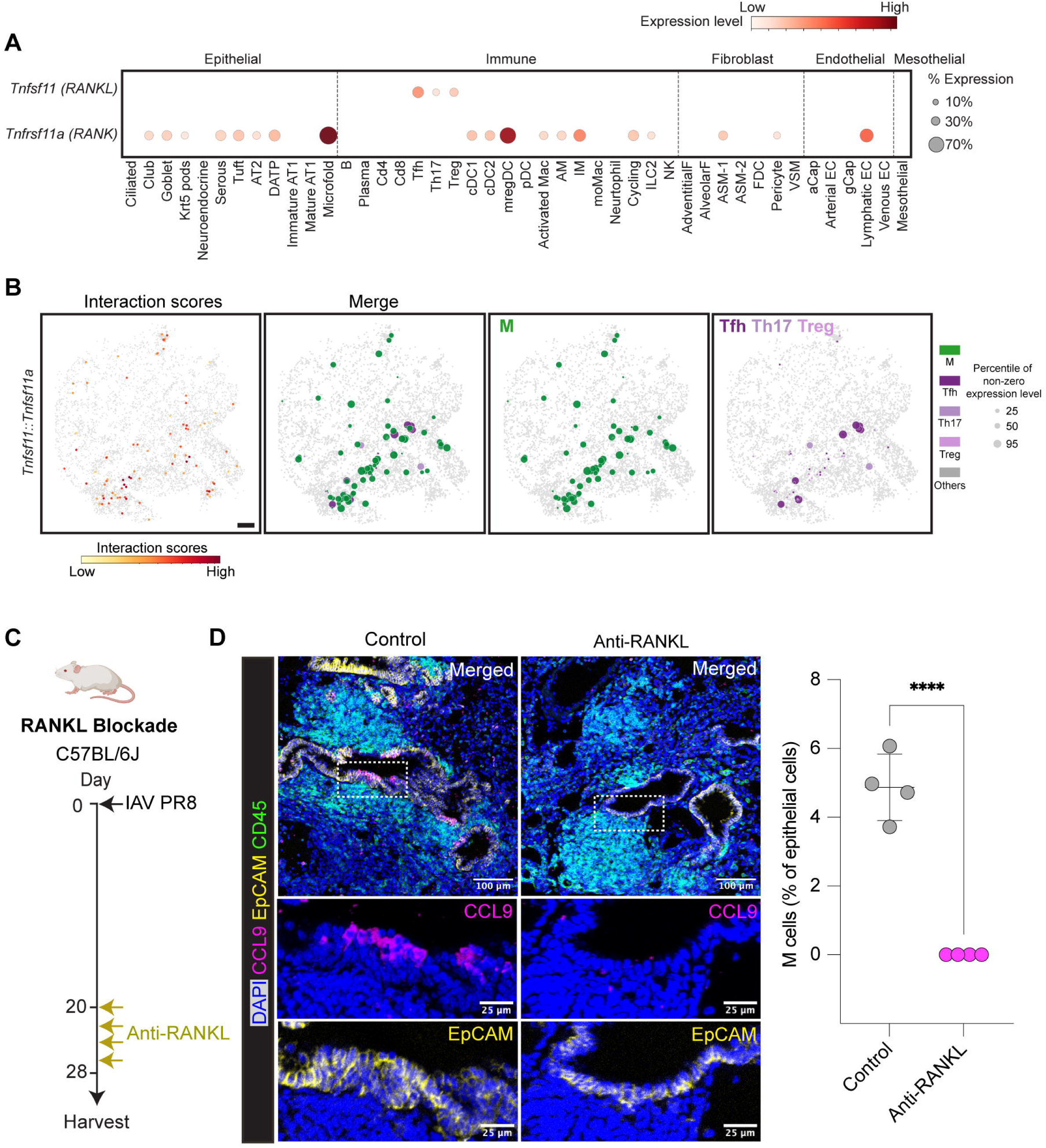
T helper cells provide RANKL signals for lung M cell differentiation. **A.** Dot plot showing expression of *RANKL* (*Tnfsf11*) and its receptor *RANK* (*Tnfrsf11a*) across all cell types. Cell types are grouped by broad compartments. **B.** Spatial distribution of RANKL-RANK interactions. The left panel shows the spatial distribution of LARIS interaction scores for RANKL-RANK interactions. The three panels on the right show RANKL-expressing immune cells and RANK-expressing M cells that constitute putative RANKL-RANK interaction pairs in a representative lung section at 4 weeks post-influenza A infection. Dot size indicates the percentile of non-zero expression level of *RANKL* in Tfh, Treg, and Th17 cells and *RANK* in M cells. Scale bars, 500 μm. **C.** Schematic of the experimental workflow for RANKL blockade in Influenza A/PR8-infected mice. **D.** (Left) Representative immunofluorescence images of airway epithelium from lung sections of wildtype mice treated with control IgG or anti-RANKL antibody. Regions outlined by the dotted squares are shown at higher magnification in the insets below. Nuclei: DAPI (blue). M cells: CCL9 (magenta), epithelium: EpCAM (yellow), immune cells: CD45 (green). Scale bars, 100 μm (main images) and 25 μm (insets). (Right) Quantification of lung CCL9-positive M cells expressed as the percentage of EpCAM positive epithelial cells, following influenza infection and RANKL blockade (n = 4 mice per group; bars indicate mean ± SD; analysis by unpaired two-tailed Student’s *t*-test; \*\*\*\**p <* 0.0001).

### Lineage tracing demonstrates that lung M cells arise from secretory progenitors that differentiate in response to RANK signaling

We next sought to define the epithelial origin of influenza-induced lung M cells. We previously demonstrated that RANKL stimulation induces M cell differentiation from both tracheal basal stem cells and secretory epithelial cells in primary epithelial cultures.^13^ Although basal stem cells constitute a major progenitor population in the trachea, they are less frequent in bronchial airways, where iBALT is known to occur. ^28,29^ However, basal cells can appear in greater numbers in bronchi following viral infection, and therefore could serve as M cell progenitors despite their paucity in the homeostatic bronchi.^29–31^ To determine the epithelial lineage of influenza-induced lung M cells *in vivo*, we performed lineage-tracing experiments using *Scgb1a1-CreER::tdTomato* and *p63-CreER::tdTomato* mice, enabling tamoxifen-induced labeling of secretory and basal stem cell lineages, respectively (Figure 3A). Lineage-labeled basal stem cells were not the progenitors for the newly differentiated lung M cell population. In contrast, lung M cells were robustly labeled by the *Scgb1a1* lineage tracing, demonstrating that influenza-induced lung M cells arise predominantly from secretory epithelial progenitors (Figure 3B).

**Figure 3.**
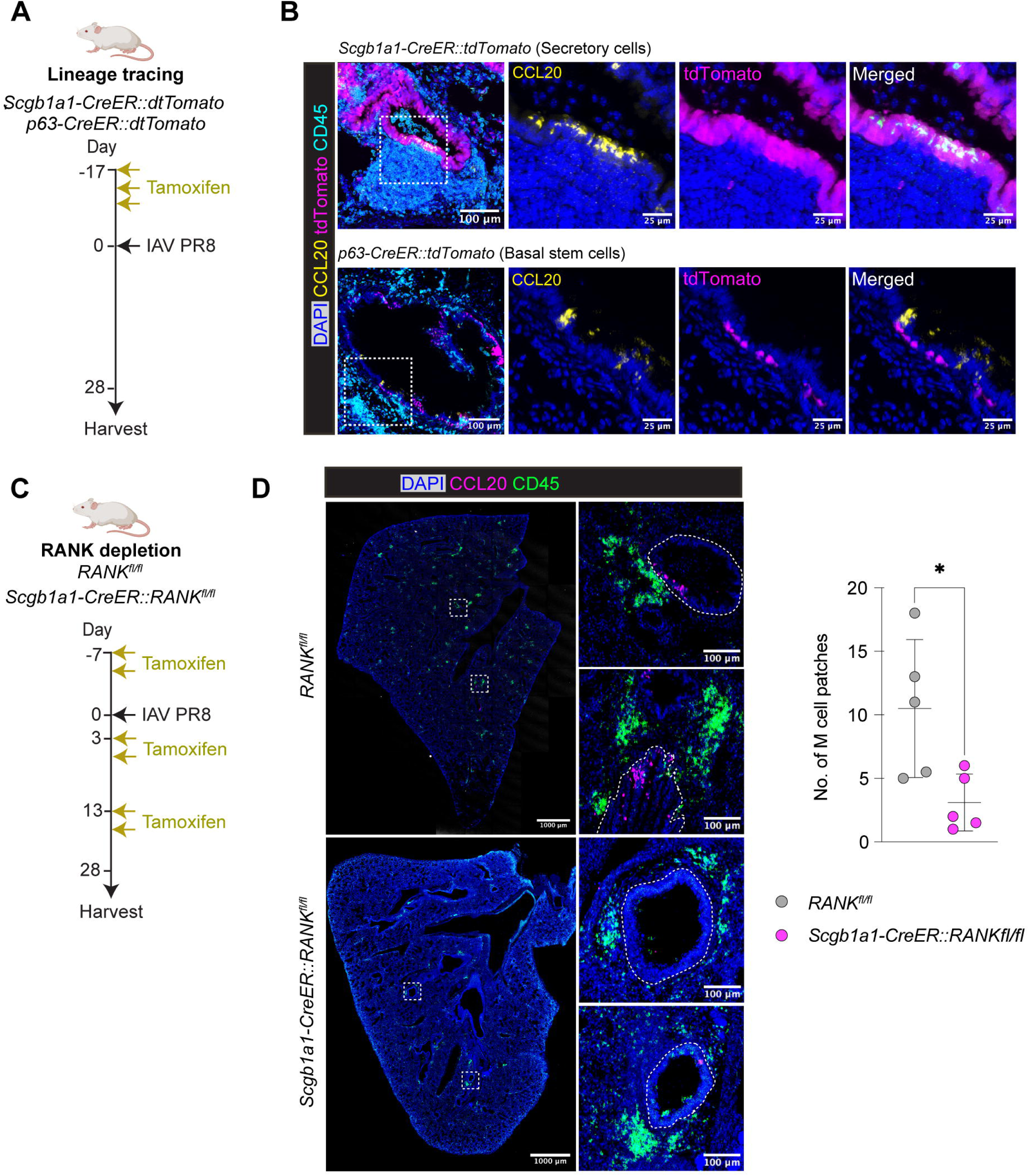
Lung M cells originate from secretory progenitors through RANK-dependent differentiation. **A.** Schematic of the experimental workflow for lineage tracing epithelial secretory and basal stem cells following Influenza A/PR8 infection. **B.** Representative confocal images of lung sections from lineage-labelled secretory cell (*Scgb1a1CreER::tdTomato*, upper panel) and basal stem cell (*p63CreER::tdTomato* mice, lower panel) lineage tracing mice following tamoxifen administration and Influenza A/PR8 infection. Nuclei: DAPI (blue), M cells: CCL20 (yellow), lineage-labelled cells: tdTomato (magenta), immune cells: CD45 (cyan). Regions outlined by the dotted squares are shown at higher magnification in the corresponding insets on the right. Scale bars, 100 μm (main images) and 25 μm (insets). **C.** Schematic workflow for tamoxifen-enabled genetic loss of RANK in secretory cells in Influenza A/PR8 infected mice. **D.** (Left) Representative immunofluorescence image of lung sections from *Rank^fl/fl^* and *Scgb1a1-CreER::Rank^fl/fl^* mice following tamoxifen treatment and influenza A/PR8 infection. The regions outlined by the dotted squares in the whole-lung sections are shown at higher magnifications in the right insets. Nuclei: DAPI (blue). M cells: CCL20 (magenta), immune cells: CD45 (green). Scale bars, 1,000 µm (whole-lung section) and 100 µm (inset). (Right) Quantification of CCL20 positive M cell patches in mouse lungs sections following influenza A/PR8 infection. Quantifications were averaged across two representative sections per mouse. (n = 5 mice per group; bars indicate mean ± SD; analysis by two-tailed Mann-Whitney U test; **\****p* < 0.05).

Having established that RANKL–RANK signaling is required for influenza-induced M cell differentiation and furthermore that lung M cells arise from secretory epithelial cells, we next asked whether secretory cell RANK-mediated signaling reception is required for their differentiation into M cells. We conditionally deleted *RANK* from secretory epithelial cells using *Scgb1a1-CreER::RANK^fl/fl^* mice (Figure 3C). Indeed, tamoxifen-induced deletion of *RANK* significantly reduced the abundance of lung M cell patches following influenza infection (Figure 3D).

Unlike the constitutively present M cells associated with secondary lymphoid organs in the gut, airway M cells are very rare under physiologic conditions. Instead, we demonstrate that lung M cells are predominantly induced alongside iBALT following influenza infection through inflammation-driven RANKL–RANK signaling. We therefore designate them as inducible microfold cells (iM cells) hereafter.

### iM cells are required for the proper organization of iBALT

Given the close spatial association and coordinated kinetics of the appearance of iM cells and iBALT following influenza infection, we next assessed whether iM cells are necessary for the proper organization of influenza-induced BALT. We employed *Sox8-CreER::DTA* mice to ablate M cells. *Sox8-CreER::DTA* mice were infected intranasally with influenza A virus/PR8, and multiple doses of tamoxifen were administered beginning on day 9 post-infection to effect continued DTA-induced M cell ablation (Figure 4A). Immunofluorescence staining of day 28 post-infection lungs revealed a tenfold depletion of CCL9⁺ iM cells (Figure 4B).

**Figure 4.**
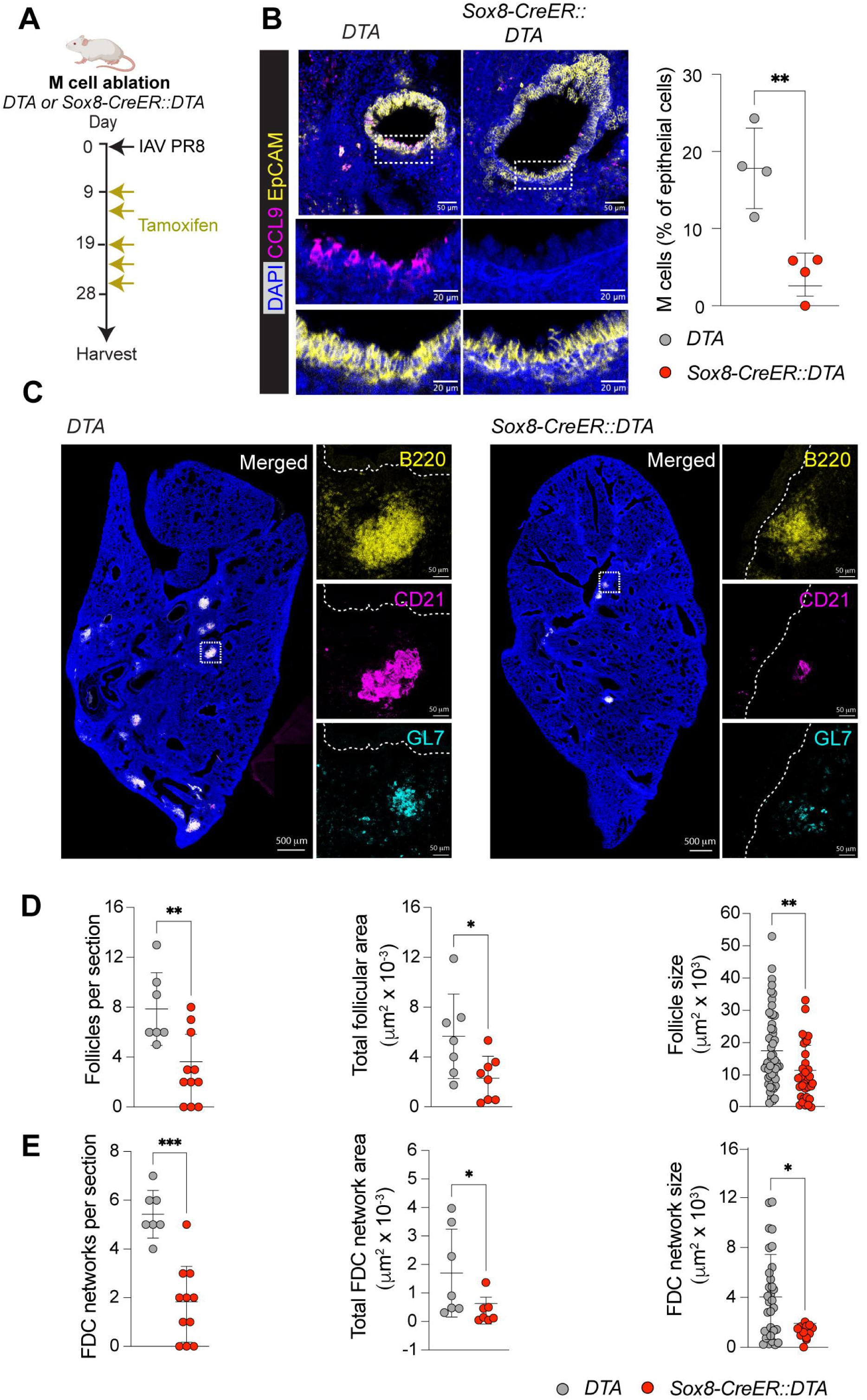
iM cell ablation reduces iBALT abundance and disrupts follicular architecture. **A.** Schematic of the experimental workflow for iM cell ablation following Influenza A/PR8 infection. **B.** (Left) Representative confocal images of lung sections from *Sox8-CreER::DTA* and DTA control mice. M cells: CCL9 (magenta). Epithelial cells: EpCAM (cyan). Scale bars, 50 µm (main images) and 25 µm (inset), (Right) Quantification of iM cells expressed as a percentage of epithelial cells, following influenza infection and *Sox8*-dependent ablation (n = 4 mice per group; bars indicate mean ± SD; analysis by unpaired two-tailed Student’s *t*-test; **\*\****p* < 0.01). **C.** Representative immunofluorescence images of iBALT structures in *Sox8-CreER::DTA* mice and DTA control littermates. Insets are marked with a dotted square in the whole lung sections. B cell follicles: B220 (yellow). Follicular dendritic cell networks: CD21 (magenta). Germinal centers: GL7 (cyan). Scale bars, 500 μm (whole-lungs sections) and 50 μm (insets). **(D-E)** Quantifications of B cell follicles (B220) (D) and FDC network (CD21) counts, total area (normalized to the area of lungs), and size distribution in *Sox8-CreER::DTA* mice compared with DTA controls. Each dot represents one iBALT structure (size measurements) or one lung section (counts and total area). (n = 7-11 mice per group, bars indicate mean ± SD, analysis by Welch’s t-test. \**p* < 0.05, ** *p* < 0.01, *** *p* < 0.001; ns, not significant).

Next, we investigated the impact of iM cell depletion on iBALT architecture. Lung sections were stained for B220, CD21, and GL7 to visualize B cell follicles, follicular dendritic cell (FDC) networks, and germinal centers (GCs), respectively. Whereas control mice developed abundant, mature iBALT structures containing compact GL7⁺ GCs supported by extensive CD21⁺ FDC networks, M cell-depleted mice exhibited fewer and smaller iBALT structures with markedly contracted FDC networks and spatially dispersed, fragmented GL7⁺ germinal centers (Figure 4C). Furthermore, iM cell ablation significantly decreased iBALT frequency. The remaining follicles displayed reduced B cell follicle size and reduced total follicular area (Figure 4D). FDC organization was disrupted and characterized by significant reductions in the number, total area, and average size of FDC networks (Figure 4E). Thus, we show that iM cells serve a critical role in maintaining proper iBALT architecture, which is known to be essential for effective mucosal immunity.^8^

### Spatial BCR profiling demonstrates that iM cell-associated iBALT supports B cell clonal expansion and diversification

A defining feature of mature, functional TLOs is their capacity to support local germinal center reactions, including B cell activation, clonal expansion, class switching, and somatic hypermutation (SHM).^3,9^ By integrating Slide-tags spatial transcriptomics with single-cell BCR V(D)J sequencing, we were able to concomitantly examine B cell transcriptional states and establish spatially delimited clonal relationships within iM cell-associated iBALT.

Combining all B cells and plasma cells identified in our Slide-tags dataset, we performed higher-resolution clustering to resolve distinct B cell states. This analysis identified naïve B cells, activated B cells, tissue-resident memory B cells (BRM), germinal center (GC) B cells, B-1 like cells, and plasma cells (Figures 5A and S3A). GC B cells were characterized by the expression of canonical germinal center-associated genes, including *Aicda, Bcl6, Cxcr4, Fas, and Mki67* (Figure S3A). Consistent with ongoing germinal center activity and immunoglobulin class-switching, naïve, activated, and GC B cell populations exhibited expression of IgM, IgA, and IgG, whereas BRM and plasma cells predominantly expressed IgA and IgG (Figures 5A and S3A).^32^

**Figure 5.**
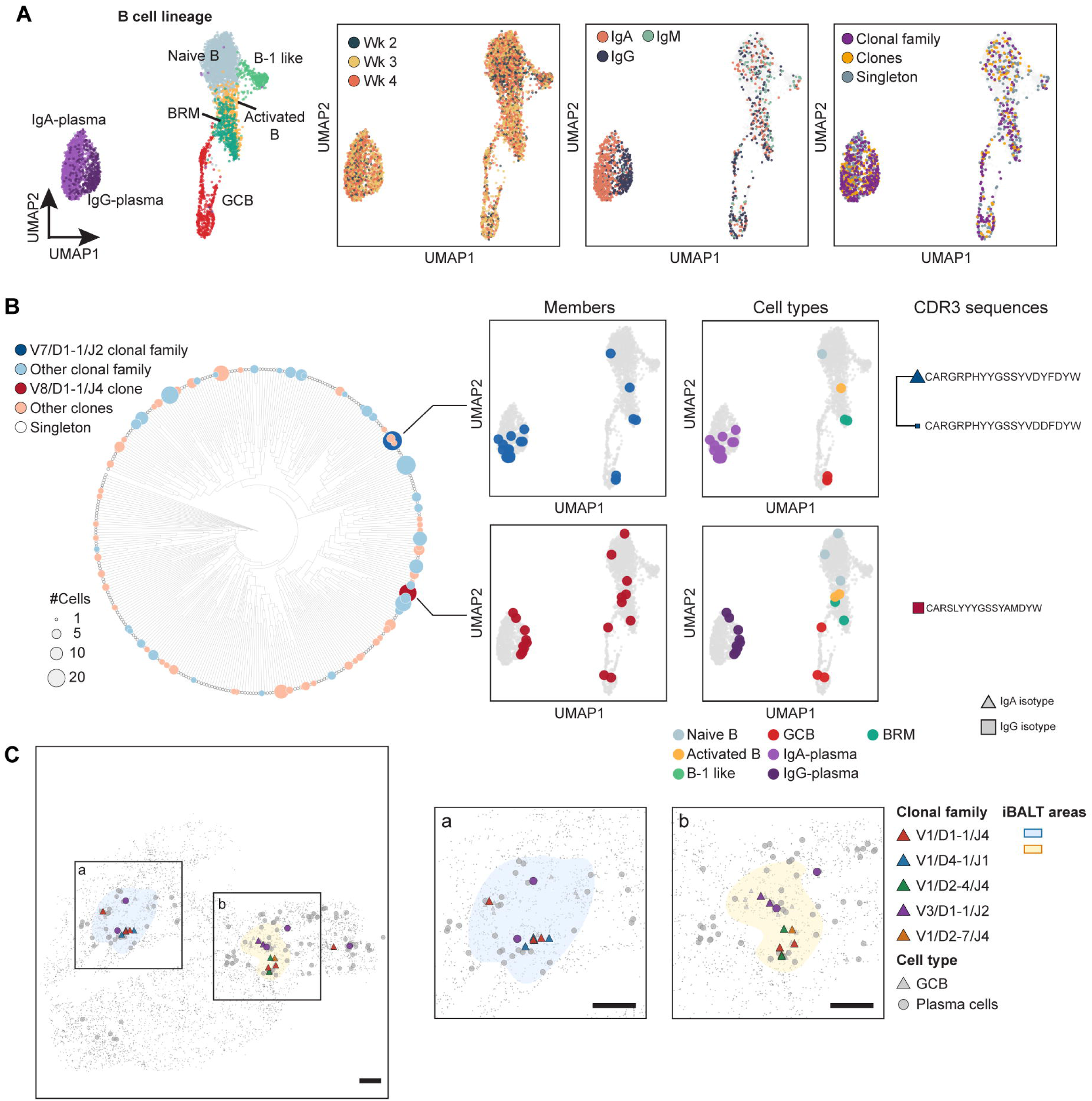
iM cell-associated iBALT structures support local B cell clonal expansion and diversification. **A.** UMAP representation of B cells at different time points post-influenza A/PR8 infection. Left panel: Cell type annotation. Mid-left panel: Timepoint of origin. Mid-right panel: BCR isotypes. BCR isotype was assigned from VDJ sequencing. Right panel: Clonal expansion status. Cells without a detected BCR are shown in light grey. **B.** Circular phylogenetic tree of BCR CDR3 sequences from the week 3.1 sample. Branch lengths reflect relative CDR3 sequence divergence. Tip color identifies clonal families (red) or clones (blue). Tip size is proportional to the number of cells in the clonal family or clone. Selected clonal families or clones are shown on the right, together with their corresponding CDR3 amino acid sequences and antibody isotypes. **C.** Spatial distribution of clonally expanded germinal center (GC) B and plasma cells across iBALT structures. Left, overview of the full tissue section, with individual iBALT structures outlined in distinct colors. GCB cells are shown as triangles and plasma cells as circles. Cells belonging to expanded clonal families containing ≥2 GCB cells within an iBALT are colored by clonal family; other GCB cells are shown in light grey and other plasma cells in grey. Right, enlarged views of representative iBALT structures (iBALT-a and iBALT-b), showing cells within and immediately surrounding each structure using the same color scheme.

We recovered BCR sequences from approximately 1,362 of 4,880 (27.9%) B cell and plasma cell nuclei (Figure 5A), enabling analysis of B cell clonal relationships. Analysis of the recovered BCR sequences revealed substantial clonal expansion and sequence diversification. We distinguished two levels of B cell clonal relatedness (Figures 5A and 5B, left panel). First, we identified expanded clones sharing identical V, D, and J gene usage and identical CDR3 sequences. Second, we identified broader groups of B cells sharing the same V(D)J gene usage and CDR3 length but containing closely related CDR3 sequences (≤20% Levenshtein distance), which we termed “clonal families”. This sequence relatedness is consistent with diversification of a common ancestral B cell clone through somatic hypermutation.^33^ The remaining BCR sequences were represented as singletons (Figures 5A and 5B, left panel, Table S1).

Both clones and clonal families spanned multiple B-cell states, including naïve, activated, GC B cells, BRMs, and plasma cells, revealing clonal relationships across distinct B cell differentiation states within iM cell-associated iBALT (Figures 5A and 5B, middle panel). A subset of clonal families contained members expressing divergent immunoglobulin isotypes, suggesting that class-switch recombination (CSR) can occur along distinct branches of a shared clonal lineage (Figure 5B, top right panel). To further characterize isotype diversity within clonally related B cells, we focused on plasma cells, as their fixed CDR3 sequences and committed isotypes make them the cleanest readout of GC output. In contrast to 100% pure isotype in clones (N = 62), 20% of the clonal family (N = 64) plasma cells showed mixed IgG and IgA isotypes (Figures S3B-3E). This suggests that, although isotype identity is highly conserved within identical-CDR3 clones, isotype diversification can occur among more distantly related members of the same clonal family. Consistent with this observation, pairwise comparisons of plasma cells within clonal families revealed that isotype concordance was significantly above chance for highly similar CDR3 sequences (distance 0-0.05; p < 0.001) but progressively declined with increasing CDR3 sequence divergence (distance 0.15–0.20; p = 0.150). Overall, CDR3 sequence divergence was negatively correlated with isotype concordance (Spearman’s ρ = −0.45; Figure S3F). B-1-like cells were largely excluded from these relationships consistent with their distinct innate-like B-cell identity (Figure 5A, Table S1).^34^ Together, these findings suggest that class switching and BCR sequence diversification can occur at different stages of B-cell clonal evolution, giving rise to distinct isotype-defined populations within shared clonal lineages.

Spatial mapping of clonal families further revealed that GC B cells belonging to the same clonal family could be found across spatially separated iBALT structures (Figure 5C), suggesting potential migration and reseeding between distinct iBALT sites or from lymph nodes. Furthermore, GC B cells belonging to multiple distinct lineages frequently coexisted within the same iM cell-associated iBALT, indicating multiclonal GC responses (Figure 5C).^35^ Together, these findings provide a spatially resolved view of B cell clonal architecture within influenza-induced BALT and reveal that iM cell-associated iBALT harbors clonally expanded and diversified B cells.

### Spatial transcriptomics reveals cytokine signaling networks within iM cell–iBALT units

Having established that iM cells are required for iBALT organization and local adaptive immune responses, we next sought to identify molecular signals through which iM cells communicate with adjacent immune cell populations. To do so, we employed LARIS,^36^ a computational framework that incorporates spatial proximity analysis to infer ligand-receptor interactions, in order to identify spatially constrained cytokine interactions between iM cells and neighboring immune populations.^36^ Using LARIS analysis, we identified multiple cytokine interactions that link iM cells with iBALT-associated B cells, T cells, and dendritic cells (Figure 6A, Table S2). Among iM cell-derived signals, the CCL20–CCR6 axis emerged as a prominent interaction linking M cells to B cells and Th17 cells. Spatial mapping of *Ccl20-Ccr6* interaction scores demonstrated that *Ccl20*-expressing iM cells localized within regions containing *Ccr6*-expressing B cells and Th17 cells (Figure 6B, left panel), identifying these populations as candidate recipients of iM cell-derived CCL20 signaling. This is consistent with the known role of CCL20–CCR6 signaling in lymphocyte recruitment and positioning within mucosal lymphoid tissues, most interestingly in the localization of memory B cells following influenza infection.^37,38^

**Figure 6.**
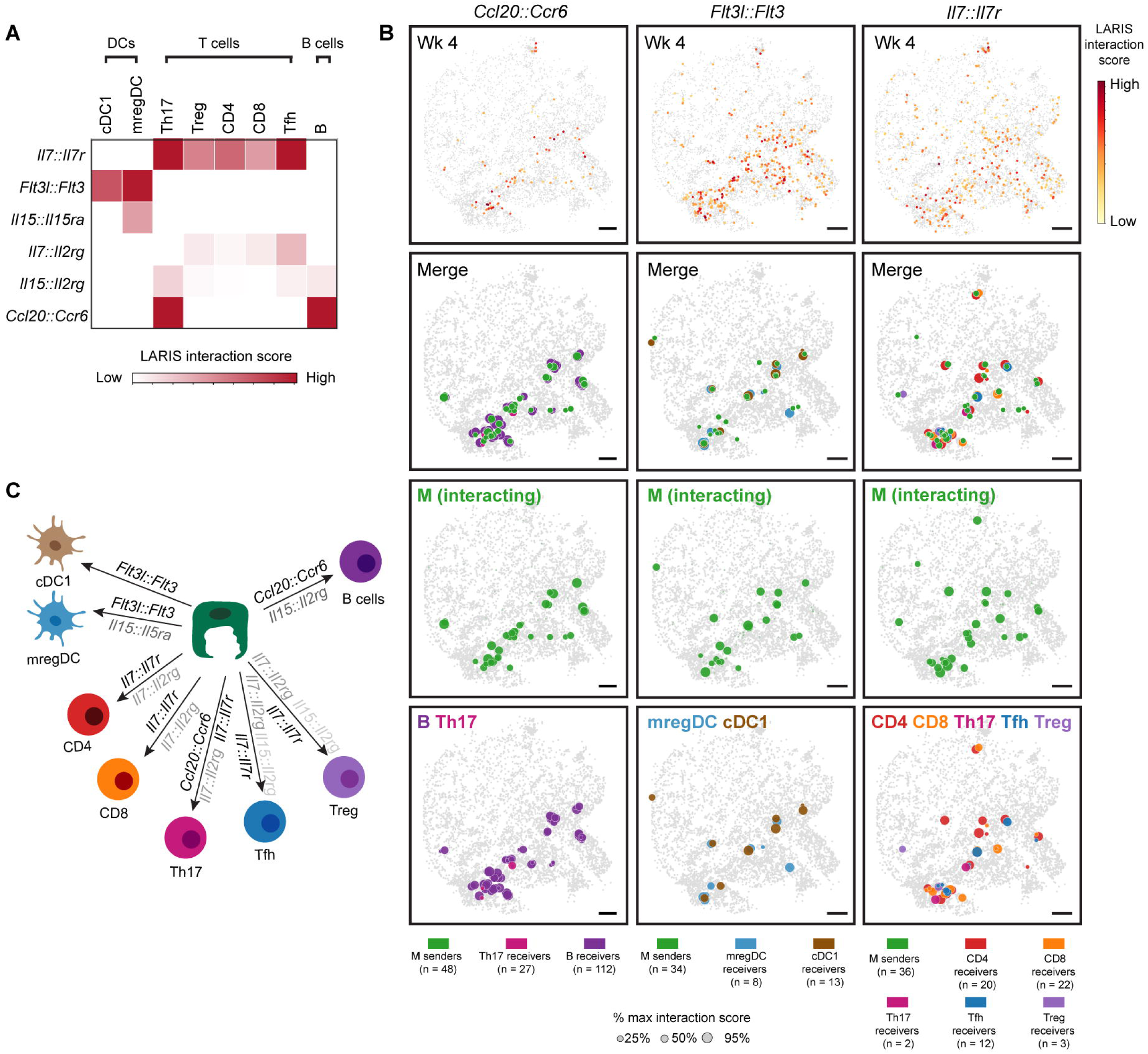
Spatial transcriptomics identifies specific iM cell to immune cell cytokine signaling networks. **A.** Ligand-receptor interactions between iM cells and neighboring immune cell types. Color intensity indicates summed LARIS interaction scores, with only interactions at FDR < 0.05 displayed. **B.** Spatial distribution of selected interactions and the corresponding immune and iM cells participating in putative interactions. The top panels show the spatial distribution of LARIS interaction scores for *Ccl20-Ccr6, Flt3l-Flt3,* and *Il7-Il7r* interactions. The three panels below show *Ccl20*-, *Flt3l*, or I*l7*-expressing iM cells and *Ccr6*-, *Flt3*-, or *Il7r*-expressing immune cells that constitute putative interaction pairs in a representative lung section at 4 weeks post-influenza infection. Dot size indicates sender scores in iM cells and receiver scores in corresponding immune cells. Scale bars, 500 μm. **C.** Summary schematic of iM-immune cell ligand-receptor interactions. Interactions are shown in varying color intensities, with darker colors indicating stronger interactions.

We also identified FLT3L–FLT3 signaling operating between iM cells and adjacent dendritic cell populations. This coincides with the observation that stromal cell-derived FLT3L supports the development and homeostatic maintenance of dendritic cells within SLOs, while FLT3L-responsive dendritic cell populations have also been implicated in the organization and maintenance of tumor-associated TLOs.^39,40^ Notably, FLT3L–FLT3 interactions were also predicted between follicular dendritic cells (FDCs) and dendritic-cell populations, with higher interaction scores observed for FDC–mregDC and FDC–cDC1 interactions than for the corresponding iM-cell interactions (Figure S4). Additionally, spatial mapping identified *Flt3l*-expressing iM cells in regions containing FLT3-expressing presumptive mregDCs and conventional type 1 dendritic cells (DC1s) (Figure 6B, middle panel), suggesting that iM cells may provide an additional epithelial source of signals supporting local dendritic-cell populations.

Lastly, LARIS analysis revealed widespread IL-7–IL-7R signaling interactions between iM cells and multiple T cell populations, including CD4 T cells, Th17 cells, Tfh cells, Treg cells, and CD8 T cells (Figure 6B, right panel). IL-7–IL-7R interactions also occurred between lymphatic endothelial cells (LECs) and the same T-cell populations, although these interactions were characterized by substantially lower interaction scores than those originating from iM cells (Figure S4). Together, these findings suggest that iM cells may represent a more prominent source of IL-7 signaling than LECs within influenza-induced pulmonary lymphoid niches, potentially providing survival and maintenance cues to neighboring T cell populations. This contrasts with previous findings implicating LEC-derived IL-7 in T cell maintenance within allergen-induced BALT, raising the possibility that the cellular sources of IL-7 and their relative contributions to lymphoid niche organization may differ depending on the inflammatory context. ^41^

In aggregate, our findings identify a network of candidate iM cell–immune cell interactions that contribute to follicular lymphocyte positioning, dendritic-cell behavior, and T cell survival, highlighting a potential role for iM cells as epithelial signaling hubs that orchestrate the organization and maintenance of iBALT (Figure 6C).

### iM cell ablation results in a loss of influenza antigen-specific germinal center (GC) B cells, T follicular helper (Tfh), and tissue-resident memory T helper (TRM) cells

We next assessed whether M cell ablation broadly alters local and peripheral immune cell composition and antigen-specific immune responses. We therefore performed flow cytometric profiling of influenza nucleoprotein (NP)-specific B and T cells from *Sox8-CreER::DTA* mice at 28 days following influenza infection. Mice were treated with anti-CD45 antibody intravenously immediately prior to tissue collection to discriminate antibody-labeled intravascular immune cells from unlabeled tissue-localized (IV⁻) immune cells.

We measured antigen-specific B cell populations using an NP protein probe (Figure S5A).^9,42^ M cell ablation markedly reduced the number of NP-specific lung germinal center (GC) B cells, whereas NP-negative GC B-cell numbers were not significantly altered (Figures 7A and S5B). This selective reduction in the NP-specific GC compartment indicates that iM cells specifically contribute to the generation and/or maintenance of antigen-specific GC B cell responses. In contrast, both the NP-specific and NP-negative lung-resident memory B (BRM) cell compartments were preserved (Figures 7B and S5C). Moreover, the abundance of NP-specific and NP-negative total B cells in the lung remained unchanged following M-cell ablation (Figure S5D). Similarly, NP-specific total B cells, GC B cells, and BRM cell numbers were unchanged in mediastinal lymph nodes (Figure S5E). Thus, the loss of iM cells selectively impaired local antigen-specific GC B cell responses.

**Figure 7.**
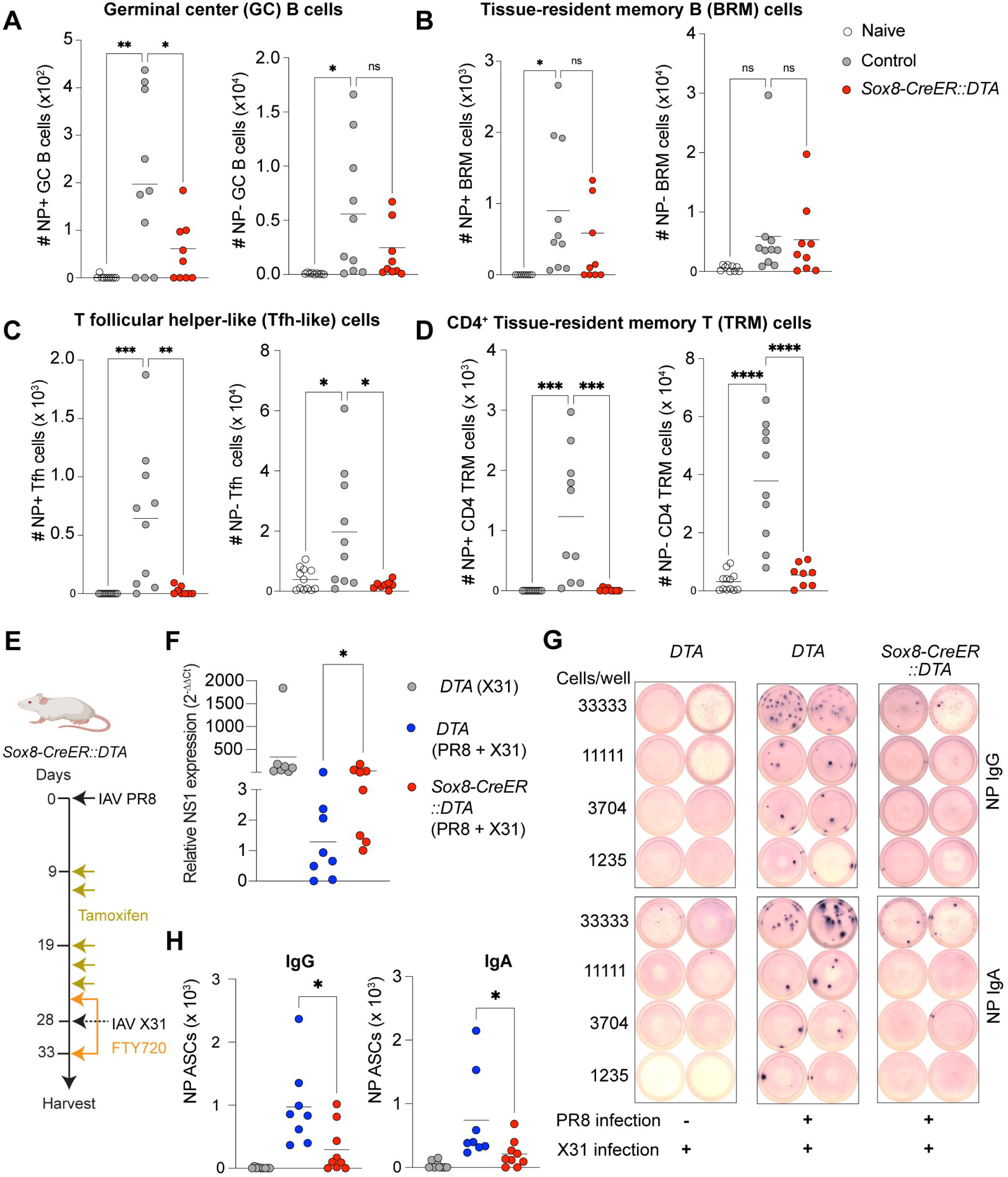
iM cells promote antigen-specific local germinal center responses and pulmonary mucosal protection following influenza rechallenge. **A-D.** Flow cytometric quantification of influenza A/PR8 nucleoprotein (NP)-specific and NP-negative immune cell populations in the lungs of control (*DTA*) and M cell-ablated (*Sox8-CreER::DTA*) mice following influenza infection. NP-specific B cells were identified using a fluorescently labeled recombinant NP protein probe, whereas NP-specific T cells were identified using NP peptide–MHC II tetramers. (A) Germinal center (GC) B cells. (B) Tissue-resident memory B cells (BRMs). (C) T follicular helper-like (Tfh-like) cells. (D) CD4^+^ tissue-resident memory T cells (TRM). PBS-treated naïve mice served as negative controls. Each symbol represents an individual mouse. Data are pooled from two independent experiments (n = 8–10 mice per group). Horizontal bars indicate the mean. Statistical significance was determined using one-way ANOVA with Tukey’s multiple comparisons test. *p < 0.05, \*\**p* < 0.01, \*\*\**p* < 0.001, \*\*\*\**p* < 0.0001; ns, not significant. **E.** Schematic for the experimental workflow for heterotypic influenza rechallenge model. **F.** Lung viral burden following heterotypic influenza A/X-31 rechallenge, assessed by quantitative PCR (qPCR) for viral NS1 RNA. Data are pooled from two independent experiments (n = 7–8 mice per group). Horizontal bar represents the mean. Statistical significance was determined using Mann– Whitney U test. \**p* < 0.05. **G.** Representative ELISpot images showing influenza NP-specific antibody-secreting cells (ASCs) from the lungs of control (*DTA*) and M cell-ablated (*Sox8-CreER::DTA*) mice following primary influenza A/PR8 and subsequent heterotypic influenza A/X-31 virus rechallenge. **H.** Quantification of NP-specific IgG- and IgA-secreting ASCs in the lungs of control (*DTA*) and M cell-ablated (*Sox8-CreER::DTA*) mice following heterotypic influenza rechallenge. Data are pooled from two independent experiments (n = 8–9 mice per group). Horizontal bars represent the mean. Statistical significance was determined using Mann–Whitney U test. \**p* < 0.05.

We next assessed the abundance of influenza NP-specific CD4 T cell populations using an NP MHCII tetramer (Figure S6A).^43^ M cell ablation significantly reduced NP-specific lung T resident helper/T follicular helper-like (Tfh-like) cells (Figures 7C and S6B). NP-negative Tfh-like cells were also reduced, indicating a broader disruption of the local follicular helper compartment (Figures 7C and S6B). Similarly, both NP-specific and NP-negative CD4 lung-resident memory (TRM) cells were markedly lowered following M cell ablation (Figure 7D and S6C). By contrast, NP-specific and NP-negative total CD4 T-cell and Treg populations in the lung did not show significant differences between groups (Figures S6D and 6E). These findings indicate that iM cells preferentially support local T follicular helper-like (Tfh-like) cells and tissue-resident memory (TRM) cells.

### iM cells are essential for mucosal IgA- and IgG-mediated protection in the setting of influenza rechallenge

To determine the functional relevance of iM cells, we performed M cell ablation in the setting of influenza rechallenge. Four weeks after primary infection with the influenza A/PR8 strain, mice were rechallenged with the heterotypic influenza A/X-31 strain.^24,42^ To restrict our analysis of the recall response to tissue-resident immune cell populations, we treated mice with FTY720, a sphingosine-1-phosphate receptor antagonist that prevents lymphocyte egress from lymphoid tissues, thereby limiting the contribution of recirculating lymphocytes to the pulmonary recall response (Figure 7E).^24,42^ M cell-ablated mice exhibited markedly higher lung viral burden, as determined by quantitative RT-PCR measurement of influenza A NS1 RNA (Figure 7F). Furthermore, NP-specific IgG and IgA lung antibody-secreting cells (ASCs) were significantly reduced (Figures 7G and H). Together, these findings demonstrate that iM cells are required for efficient local humoral recall and viral control during heterotypic influenza rechallenge.

## Discussion

Barrier epithelia are increasingly appreciated as active participants in tissue immunity.^44,45^ In many mucosal organs, specialized regions of the epithelium are coupled to immune follicles. These composite structures form functional lymphoid organs that support mucosal immunity.^10,20^ In so-called secondary lymphoid organs (SLOs), these structures are embryonically patterned. In contrast, tertiary lymphoid organs (TLOs) are thought to form in response to infection and inflammation, and notably in the setting of cancer. Despite the recent flourish of interest in TLOs, their formation and function are only beginning to be understood.^1,46^ Here, we define the epithelial and immune origins of lung TLOs. While the existence of inducible bronchus-associated lymphoid tissue (iBALT) has long been recognized, we now show for the first time that iBALT is functionally linked to inducible microfold (iM) cells of the airway epithelium, and these iM-iBALT units function as TLOs that support local adaptive immunity to lung infection.

We and others have previously shown that airway M cells take up antigen and infectious particles.^11,13,26,47,48^ Here, we show that iM cell ablation results in disrupted iBALT architecture and an accompanying reduction in the abundance of antigen-specific germinal center B cells, T follicular helper cells, and importantly, tissue-resident memory CD4^+^ T cells. These changes are accompanied by a decrement in influenza-specific mucosal IgA and IgG responses and ultimately lead to compromised viral control. Recent elegant work has shown that M cells in intestinal SLOs not only process antigens, but also directly present these antigens.^49^ It will be intriguing to establish whether such is also the case in TLOs in general and more specifically in lung TLOs. We speculate that the instability of iM cells may limit TLO lifespan in contrast to the ever-present SLOs of the gut, where M cells are a persistent feature of the epithelium. Indeed, one wonders whether the lifespan of iM cells will vary according to inflammatory context. Finally, since it is known that the gut can develop TLOs in response to inflammation and infection, presumably to amplify SLO function, it will be of interest to establish whether these gut TLOs form in the same manner as lung TLOs.^46^

Current models of TLO formation are largely based on analogy to the mechanisms established in gut SLOs and lymph nodes.^2,46^ At the outset of embryonic follicle formation in the gut, a subset of precursor mesenchymal cells is patterned by embryonic day 15.5 (E15.5), and these cells will eventually mature into functional lymphoid tissue organizer (LTo) cells. The differentiation of LTo cells is governed by fetal liver-derived Id2^+^RORγt^+^IL-7Rα^+^CD3^-^ lymphoid tissue inducer (LTi) cells that home to the location of the aforementioned mesenchymal LTo-precursor cells through unknown mechanisms.^50–53^ The maturation of precursor mesenchymal cells into LTo cells is mediated by LTi cell-derived lymphotoxin signaling. Interestingly, lymphotoxin signaling, presumably also originating from LTi cells, is required for the specialization of the nearby intestinal epithelium into the follicle-associated epithelium (FAE). After their maturation, LTo cells serve as chemokine rich hubs that recruit lymphocytes into forming follicles.^2,46^ Subsequently, mesenchymal-derived M cell inducer cells (MCi), located in the subepithelial dome sandwiched between the FAE and the adjacent follicle, employ a membrane-bound RANKL signal that leads to the differentiation of M cells within the FAE, representing the final step in the generation of a functional SLO.^14^ The patterning of MCi cells is not understood; however, the simplest model would place their positioning downstream of the positioning of LTo precursor cells. If so, these mesenchymal precursor cells, prior to their maturation into LTo cells, would be the earliest known organizers of gut SLOs. However, even in the case of the gut, the embryonic patterning of the pre-LTo mesenchymal organizer cells remains mysterious.

Unlike the case of embryonically patterned SLOs, the precise positioning of TLOs in the adult airway is difficult to predict. In fact, LTi cells are not needed during iBALT formation, suggesting that TLOs are patterned by distinct mechanisms.^7^ Moreover, infiltrating inflammatory CD4^+^ T cells are known to employ IL-17 signaling in order to activate LTo-like programs including CXCL13, CCL19, and CCL21 in lung tissue-resident mesenchymal precursor cells, suggesting a deep homology of aspects of follicle formation.^7,54^ However, no correlate of a RANKL-expressing MCi cell has been established in lung TLOs. We now find that infiltrating Th17, Tfh, and Treg cells are the key sources of RANKL in the lung. Thus far, we have not established whether secreted or cell-surface-localized RANKL dictates the site of M cell differentiation,^14^ nor do we know whether iBALT is associated with a specialized patch of epithelium analogous to the intestinal FAE.^55^

We propose a theoretical model in which our general inability to predict the precise localization of lung TLOs is due to the fact that these inducible lymphoid organs are induced by infiltrating lymphocytes whose location varies according to the patchy inflammation that is characteristic of lung diseases.^16,17^ It would be interesting to determine whether there are early patchy IL-17-expressing immune cell infiltrates or other patchy infiltrating immune cells that cause the differentiation of a population of putative mesenchymal pre-LTo cells whose origins are unknown even in the deeply scrutinized intestinal system.^7^ Indeed, whether in lymph nodes, SLOs, or TLOs, the earliest steps ultimately presaging the site of eventual follicle formation remain open areas of inquiry.

As to the function of iM cells in the lung, we show that these iM cells serve as cytokine-producing epithelial hubs that influences the positioning and maintenance of multiple cellular components of the iBALT niche, suggesting the possibility of an analogous function for gut M cells. In our case, LARIS analysis allowed us to nominate candidate cytokine interactions occurring between iM cells and follicle-associated lymphocytes and dendritic cells.^36^ CCL20–CCR6 signaling linked iM cells with B cells and Th17 cells. FLT3L–FLT3 signaling linked iM cells with mregDCs and cDC1s. And finally, IL-7–IL-7R broadly linked iM cells with multiple T cell populations (CD4, CD8, Tfh, Th17, and Tregs). Recent elegant work in Peyer’s patches has similarly identified CCL20–CCR6 signaling from M cells to ILC3s as necessary for ILC3 localization.^56^ Finally, consistent with our findings employing cell-type-specific genetic M cell ablation, RANK deletion from intestinal epithelial cells results in a complete loss of M cells and associated follicle immaturity, alongside an attendant compromise in mucosal IgA responses.^10,25^

Functionally, we show that lymphocytes within iM cell-iBALT structures express *Aicda, thereby* establishing a mechanistic basis for somatic hypermutation and class switching. Spatial BCR profiling provided further evidence of productive germinal center activity, revealing clonally expanded and sequence-diversified B cell populations that span the maturation of GC B cells towards tissue-resident memory B cells (BRMs) and plasma cells. Importantly, we find clones that include plasma cells that express IgA and plasma cells that express IgG. Furthermore, we detected somatically diversified members of the same B cell clones spread across spatially distinct iBALT structures, implying immune trafficking. Such clonal connectivity raises the possibility that B cells traffic from one iBALT structure to another within the lung. Alternatively, it has recently been demonstrated that lymphocytes activated in the nasal-associated lymphoid tissue (NALT) can traffic to respiratory tissues, providing an additional trafficking route that would be consistent with our observation.^57^ Thus, we propose that iM-iBALT units may also function to provide niches for cells originating in other lymphoid organs. With regard to another possible immune niche function, previous studies have shown that CCR6-expressing lung BRMs localize within iBALT and contribute to local antibody responses during influenza rechallenge.^38,58^ In this context, we speculate that M cell-derived CCL20 may facilitate the recruitment of influenza-specific CCR6 expressing BRMs within iBALT, thereby enhancing mucosal antibody production during viral rechallenge.

TLOs have been observed in chronic respiratory infections, COPD, asthma, and lung cancer. Additionally, they occur in autoimmune and fibrotic lung diseases and following lung transplantation.^59,60,61^ However, their localization, cellular composition, organization, and functional consequences vary markedly.^62–64^ TLOs in lung cancer may contribute to antitumor immunity and sometimes predict favorable responses to immune-checkpoint blockade.^1^ Conversely, TLOs can sustain autoreactive lymphocytes and local antibody responses in autoimmune and chronic inflammatory diseases.^46^ Our findings raise the possibility that organ-specific epithelial cells, possibly induced by damage or neoplasia, may shape the organization and function of TLOs. Additionally, we speculate that lung TLOs containing M cells may have relevance in lung diseases associated with chronic antigen provision, including chronic infections and autoimmune diseases, as recently demonstrated in the gut.^49,65^ Defining whether analogous epithelial–lymphoid interactions operate in diverse disease-associated TLOs may point to therapeutic opportunities for a whole host of infectious, inflammatory, and neoplastic diseases, including the possibility of enhancing mucosal vaccination and antitumor immunity.

## Acknowledgements

The *Sox8-CreER* mouse was generated by the Harvard Genome Modification Core Facility. Flow cytometric analysis was performed in the MGH’s Department of Pathology, Flow and Image Cytometry Research Core, which received support from the NIH Shared Instrumentation program with grants 1S10OD012027, 1S10OD016372, 1S10RR020936, and 1S10RR023440. Confocal microscopy was performed at the CRM Confocal Microscopy Core Facility at MGH. ELISPOT assay reading was performed in Dr. Gilles Benichou lab at MGH. We would like to thank Drs. Avinash Waghray, Jiajie Xu, Gergana Shipkovenska, Chaim Chernoff, Haoxuan Tang, and other members of the Rajagopal lab, the Vyas lab (Columbia University), and the Klein lab (University of Wisconsin– Madison) for their valuable feedback.

This work was supported by NIH R01HL164563 (J.R.), R01HL157221 (J.R.), K99HL171884 (M.S.), K99HL181185 (D.S.), R01HG010647 (F.C.), NIDCD R01DC022650 (B.L.), and American Lung Association Catalyst Award CA-1447794 (D.S.). F.C. acknowledges support from the Searle Scholars Award, the Burroughs Wellcome Fund CASI Award, the Merkin Institute, Harvard Stem Cell Institute, and the NYSCF. J.R. was a NYSCF Robertson Investigator and F.C. is a current NYSCF Robertson Investigator.

## Author contributions

M.V.S., D.S., J.W.G., F.C., and J.R. designed the study. B.L. generated the *Sox8-CreER* driver line. M.V.S., V.I.C., H.W., and J.W.G. performed all the flow cytometry studies. M.V.S., A.X., and J.V.H. performed microscopy and image quantifications. M.V.S., and V.P. performed ELISPOT assay. M.V.S., D.S., V.P., S.K., Q.L., J.V.H. J.W.G., V.I.C., H.W., J.S., S.Z., and V.S.S. conducted all the mice experiments. P.K.T. performed cryo-sectioning. D.S. performed Slide-tags spatial transcriptomics and BCR profiling. D.S. and T.T performed LARIS analysis. D.S. and J.A.W performed neighborhood enrichment analysis. G.M. and V.P. designed the pucks for Slide-tags. M.V.S and D.S. analyzed the data, interpreted the results, and designed the figures. M.V.S., D.S., and J.R drafted the manuscript. M.V.S., D.S., B.L., J.W.G., F.C., and J.R. secured the funding. All authors read and approved the final version of the manuscript.

---

Figure S1. Validation of the *Sox8-CreER* M cell driver mouse **A.** Schematic illustrating the strategy for lineage labeling of lung M cells in *Sox8-CreER::LSL-tdTomato* mice using the tamoxifen-inducible CreER-loxP system. **B.** Representative immunofluorescence image of a lung section showing lineage-labeled lung M cells. Mature M cells identified by co-expression of CCL9 and GP2 are labeled with tdTomato. The boxed region is shown at higher magnification in the insets. Scale bars, 100 μm (whole image) and 20 μm (insets). **C.** Quantification of the percentage of CCL9⁺GP2⁺ M cells labeled with tdTomato in *Sox8-CreER::LSL-tdTomato* mice. **D.** Quantification of the specificity of Sox8-CreER lineage labeling, expressed as the percentage of tdTomato⁺ cells that co-express CCL9 and GP2.

Figure S2. Spatial transcriptomics reveals distinct cellular niches in influenza-injured mouse lungs. **A.** Distribution of cells across different cellular compartments and time points. **B.** Spatial map of cell types of the tissue section profiled. Cells are colored according to cell type annotation. Scale bars denote 500 μm. **C.** Spatial co-enrichment of cell-type pairs in lung samples 2 weeks (top) or 4 weeks (bottom) post-influenza infection within a 200-µm radius. Columns denote center cell types and rows denote neighboring cell types. Colors indicate the permutation-derived co-occurrence Z-scores, and circle size represents the median number of center cells within 200 µm of each neighboring cell. Boxes highlight cell types associated with distinct spatial compartments of the lung, corresponding to the alveoli, airways, and iBALT, from left to right. **D.** Dot plot showing marker gene expression for selected immune cell types. **E.** Spatial distribution of M cells alongside B cells, FDCs or Tfh cells in representative lung tissue sections 2 weeks (left panels) or 4 weeks (right panels) post-influenza infection. Scale bars denote 500 μm.

Figure S3. Clonal diversity and isotype heterogeneity of plasma cells in influenza A injured mouse lung. **A.** Dot plot showing marker gene expression for B cell lineage subtypes. **B.** Distribution of Ig isotypes in plasma cells. **C.** Percentage of clones and clonal families classified as having pure or mixed isotypes. Pure-isotype clones or families were defined as those containing plasma cells expressing exclusively IgA, IgG (including subclasses), or IgM, whereas mixed-isotype clones or families contained plasma cells expressing more than one of these isotypes. **D.** Stacked plots showing the isotype composition of plasma cells within individual clones (top) or clonal families (bottom). Each column represents a single clone or clonal family, with different colors indicating distinct isotypes. **E.** Representative examples of clonal families exhibiting pure (left) or mixed (right) isotypes. **F.** Relationship between CDR3 sequence divergence and isotype concordance. Pairwise CDR3 nucleotide distances and isotype concordance were calculated for plasma cells within clonal families. Pairs were grouped by normalized CDR3 edit distance, with mean isotype concordance and 95% confidence intervals (red shading) shown for each bin. The dashed gray line indicates the expected concordance under random isotype assignment, while the dashed green line represents the expected concordance in the absence of subsequent isotype switching. Statistical significance was assessed using one-sided binomial tests (\*\*\**p* < 0.001, \*\**p* < 0.01, \**p* < 0.05; ns, not significant).

Figure S4. Ligand-receptor analysis identifies candidate iM cell to immune cell cytokine signaling networks Ligand-receptor interactions between multiple sender cells (iM cells, FDCs, and LECs) and neighboring immune cell types. Color intensity indicates summed LARIS interaction scores, with only interactions at FDR < 0.05 displayed.

Figure S5. Assessment of lung B cell populations in M cell-ablated mice following influenza infection **A.** Gating strategy for various B cell populations in murine lungs. **B.** Representative flow cytometry plots identifying NP-specific (NP⁺) and NP-negative (NP−) germinal center (GC) B cells in control and *Sox8-CreER::DTA* mice following influenza infection. **C.** Representative flow cytometry plots identifying NP⁺ and NP− tissue-resident memory B cells (BRMs). **D.** Flow cytometric quantification of influenza A/PR8 nucleoprotein (NP)-specific and NP-negative total B cells in the lungs of control (*DTA*) and M cell-ablated (*Sox8-CreER::DTA*) mice following influenza infection. NP-specific B cells were identified using a fluorescently labeled recombinant NP protein probe. PBS-treated naïve mice served as negative controls. Each symbol represents an individual mouse. Data are pooled from two independent experiments (n = 9–10 mice per group). Horizontal bars indicate the mean. Statistical significance was determined using one-way ANOVA with Tukey’s multiple comparisons test. ns, not significant. **E.** Flow cytometric quantification of influenza A/PR8 nucleoprotein (NP)-specific total B cells, GC B cells, and BRM cells in the mediastinal lymph nodes (mLN) of control (*DTA*) and M cell-ablated (*Sox8-CreER::DTA*) mice following influenza A/PR8 infection. n = 5–16 mice per group. Horizontal bars indicate the mean. Statistical significance was determined using one-way ANOVA with Tukey’s multiple comparisons test. ns, not significant.

Figure S6. Assessment of lung T cell populations in M cell-ablated mice following influenza infection **A.** Gating strategy for various T cell populations in murine lungs. **B.** Representative flow cytometry plots identifying NP-specific (NP⁺) and NP-negative (NP−) T follicular helper-like (Tfh-like) cells in control and *Sox8-CreER::DTA* mice following influenza infection. **C.** Representative flow cytometry plots identifying NP⁺ and NP− CD4+ tissue-resident memory T (TRM) cells. **D–E.** Flow cytometric quantification of influenza A/PR8 nucleoprotein (NP)-specific and NP-negative total T cells (D) and T regulatory cells (Tregs) (E) in the lungs of control (*DTA*) and M cell-ablated (*Sox8-CreER::DTA*) mice following influenza infection. NP-specific T cells were identified using NP peptide–MHC II tetramers. PBS-treated naïve mice served as negative controls. Each symbol represents an individual mouse. Data are pooled from two independent experiments (n = 9–10 mice per group). Horizontal bars indicate the mean. Statistical significance was determined using one-way ANOVA with Tukey’s multiple comparisons test. \**p* < 0.05; ns, not significant.

**Table S1.** B cell receptor repertoire and clonal family.

**Table S2.** Significant cell-cell interactions identified by LARIS analysis.

## STAR METHODS

### KEY RESOURCES TABLE

| REAGENT or RESOURCE | SOURCE | IDENTIFIER |
|---|---|---|
| Antibodies |  |  |
| Goat anti mouse alkaline phosphatase conjugated IgA | Southern Biotech | 1040-04 |
| Goat anti mouse alkaline phosphatase conjugated kappa antibody | Southern Biotech | 1050-04 |
| CCL9 | R&D Systems | AF463 |
| CCL20 | R&D Systems | AF760 |
| GL7 AF647 | BioLegend | 144606 |
| CD45 AF488 | BioLegend | 103122 |
| EpCAM PE | BioLegend | 118205 |
| B220 AF488 | BioLegend | 103225 |
| CD21 AF594 | BioLegend | 123426 |
| CD45.2 AF700 | BioLegend | 109822 |
| CD19 FITC | BioLegend | 152404 |
| CD38 BUV396 | BD Biosciences | 740245 |
| GL7 PE | BioLegend | 144607 |
| CXCR3 BV421 | Biolegend | 126521 |
| IgD PE-Cy7 | Invitrogen | 25-5993-82 |
| IgM PerCP-Cy5.5 | Invitrogen | 46-5790-82 |
| CD4 BV605 | BioLegend | 100451 |
| CD3 BUV396 | BD Biosciences | 569614 |
| CD69 APC | BioLegend | 104513 |
| CD44 PE | BioLegend | 103008 |
| FR4 PerCP-Cy5.5 | BioLegend | 125018 |
| Fc Block | Invitrogen | 14-0161-85 |
| *InVivo*MAb anti-mouse RANKL | BioCell | BE0191 |
| *InVivo*MAb rat IgG2a isotype control | BioCell | BE0089 |
| Influenza NP<sub>311−325</sub> peptide-loaded I-A<sup>b</sup> MHC class II tetramer | NIH Tetramer Core Facility |  |
| Bacterial and virus strains |  |  |
| Influenza virus A/PR/8/34 | ATCC | VR-1469 |
| BL21 Competent E. coli | New England Biolabs | C2530H |
| E. coli DH5α pGEX4T2 RANK-L recombinant plasmid | Addgene | 108571 |
| Chemicals, peptides, and recombinant proteins |  |  |
| Influenza Nucleoprotein / NP | Sino Biological | 11675-V08B |
| FTY720 | Sigma-Aldrich | SML0700 |
| Tamoxifen | Sigma-Aldrich | 0000539420 |
| Liberase TM | Roche | 05401127001 |
| DNaseI | Roche | 10104159001 |
| Fixable Viability Dye eFluor 780 | ThermoFisher | 65-0865-14 |
| Isopropyl β-D-1-thiogalactopyranoside (IPTG) | Millipore-Sigma | I5502 |
| Potassium sulfate | Millipore-Sigma | 223492 |
| D-(+)-Glucose | Millipore-Sigma | 49139 |
| Sodium sulfate | VWR | BDH9302 |
| HEPES | VWR | 0511 |
| Magnesium chloride | Thermo Fisher Scientific | AM9530G |
| Triton X-100 solution | Millipore-Sigma | 93443 |
| Bovine Serum Albumin | LGC Clinical Diagnostics | 1900-0016 |
| NxGen RNase Inhibitor | LGC Biosearch Technologies | 30281 |
| TRIzol | Invitrogen | 15596026 |
| RNase Zap | Invitrogen | AM9780 |
| gentleMACS ® Tissue Dissociator | Miltenyi Biotech |  |
| Chloroform | Sigma Aldrich | 366927 |
| Isopropyl alcohol | Fisher Bioreagents | BP2618-500 |
| GeneAmp ®™ 10X RT buffer and MgCl<sub>2</sub> | Applied Biosystems | N8080010 |
| dNTPs | Applied Biosystems | N8080260 |
| Oligo d(T) | Invitrogen | N8080128 |
| Random Hexamers | Invitrogen | N8080127 |
| RNaseOUT™ Ribonuclease Inhibitor | Invitrogen | 10777019 |
| MultiScribe™ Reverse Transcriptase | Applied Biosystems | 4311235 |
| FastStart SYBR® Green Master | Millipore Sigma | 04673484001 |
| TempPlate® 96-Well Semi-Skirt 0.1 mL PCR Plates | USA Scientific | 1402-9990 |
| Isoflurane | Patterson Veterinary | 07-894-9580 |
| Corn oil | Sigma-Aldrich | C8267 |
| Dimethyl sulfoxide (DMSO) | Sigma-Aldrich | D2650 |
| Ketamine | Dechra |  |
| Xylazine | Patterson Veterinary | 07-895-0792 |
| Paraformaldehyde (PFA) | SantaCruz Biotechnology | SC - 281692 |
| Sucrose | Sigma Aldrich | S7903 |
| Tween-20 | Sigma Aldrich | 655204 |
| Tissue-Tek O.C.T. Compound | Sakura Finetek | 4583 |
| Superfrost Plus microscope slides | Fisher Scientific | 1255015 |
| Critical commercial assays |  |  |
| BCIP/NBT | Vector Laboratories | SK-5400 |
| FoxP3 staining buffer set | eBioscience | 00-5523-00 |
| APC Conjugation Kit - Lightning-Link<sup>®</sup> | Abcam | ab201807 |
| GST Spin Purification Kit | Pierce, Thermo Scientific | PI16108 |
| SPRIselect Beads | Beckman Coulter | B23318 |
| Qubit dsDNA Quantification Assay Kits, High Sensitivity | Thermo Fisher Scientific | Q32853 |
| TapeStation D5000 Reagents | Agilent | 5067-5589 |
| Chromium Next GEM Single Cell 5′ Kit v2 | 10x Genomics | PN-1000263 |
| Experimental models: Organisms/strains |  |  |
| *C57BL/6J* | The Jackson Laboratory | 000664 |
| *DTA* | The Jackson Laboratory | 009669 |
| *RANK<sup>fl/fl</sup>* | The Jackson Laboratory | 027495 |
| *tdTomato* | The Jackson Laboratory | 007914 |
| Oligonucleotides |  |  |
| GAPDH -F’<br>GAPDH -R’<br>NS1-F’<br>NS1-R’ | Integrated DNA Technologies |  |
| Software and algorithms |  |  |
| Slide-tags tools | Russell et al., 2023 | GitHub repository |
| Seurat v5 | Hao et al., 2024 | Seurat website |
| MiXCR (v4.7.0) | Bolotin et al., 2015 | GitHub repository |
| LARIS (v0.12.0) | Dai et al., 2025 | GitHub repository |
| Other |  |  |
| Multiscreen IP HTS filter plates | EMD Millipore | MAIPS4W10 |
| RBC lysis buffer | E-bioscience | 00-4333-57 |
| RPMI 1640 | Sigma-Aldrich | R8758 |
| 70-µm cell strainer | Corning | 431751 |
| Carbonate buffer | ThermoFisher | J63658.AP |
| Fetal bovine serum (FBS) | Gibco | A56704-02 |
| L-glutamine | Gibco | 35050-061 |
| Penicillin–streptomycin | Gibco | 15140-122 |
| Sodium pyruvate | Gibco | 11360-070 |
| β-mercaptoethanol | Sigma Aldrich | M7522 |

### RESOURCE AVAILABILITY

#### Lead contact

Further information and requests for resources should be directed to and will be provided by the lead contact, Jayaraj Rajagopal.

## Materials availability

The *Sox8CreER* mouse is available on request.

## Data and code availability

The raw data and processed data generated in this study will be submitted to NCBI Gene Expression Omnibus. The analysis pipeline will be deposited in the GitHub repository.

## EXPERIMENTAL MODELS AND STUDY PARTICIPANT DETAILS

### Mice

Male and female mice aged 12–18 weeks were used for all experiments. The *Sox8-CreER* knock-in mouse was generated using CRISPR-Cas9 microinjection as described earlier^19^. *C57BL/6J* (stock no. 000664), *DTA* mice (stock no. 009669), *RANK^fl/fl^* mice (stock no. 027495), and *tdTomato* mice (stock no. 007914) were purchased from the Jackson laboratory*. p63-CreER*^66^ *and Scgb1a1-CreER*^67^ mice were previously described.

Mice were housed in an Association for Assessment and Accreditation of Laboratory Animal Care-accredited animal facility at the Massachusetts General Hospital (MGH). All procedures were approved by the MGH Institutional Animal Care and Use Committee (IACUC). Standard housing conditions were used throughout, unless otherwise noted (temperature of 68–79°F, humidity of 30–70%, a 12 h light–dark cycle, and regular rodent diet).

### Viruses

Influenza strains PR8 was procured from ATCC (VR-1469). Influenza A/X-31 was obtained from Dr. Jason W. Griffith (MGH)

## METHOD DETAILS

### Recombinant mouse RANKL expression and purification

pGEX4T2 RANK-L recombinant plasmid (Addgene) was transformed into the BL21 competent *Escherichia coli* cells (New England Biolabs). The cultures were induced with 100 μM IPTG for 16 h at 20°C and GST-RANKL was purified from bacterial lysate by affinity chromatography using a GST Spin Purification Kit (Pierce, Thermo Scientific). Recombinant GST used as a control was prepared by the same method using empty pGEX4T2 (obtained from Dr. Zhixun Dou, MGH).

### In vivo models

Influenza PR8 was intranasally administered at 0.5 x 10^6^ CEID_50_/ml for all primary infections. Influenza X31 was used at 5 x 10^5^ EID_50_ for secondary infections. Mice were anesthetized with vaporized isoflurane and infected intranasally with 20-to 30-μl volume of virus diluted in PBS. Tamoxifen (Sigma-Aldrich) was dissolved in corn oil (20 mg/ml) and stored at 4 °C for the duration of treatment. Mice were treated with multiple doses of tamoxifen (100 mg/kg/dose) via oral gavage. For ectopic induction of M cells, GST-RANKL fusion protein was administered as daily intraperitoneal injections of 100 μg/day for 6 days. Recombinant GST prepared from empty pGEX4T2 vector was used as a control for GST-RANKL. Mice were treated with anti-RANKL antibody (BioCell) intraperitoneally for *in vivo* RANKL neutralization experiments as described previously.^25^ Mice were treated with 250 μg of purified anti-RANKL or a control rat IgG2a antibody on four alternate days before sacrifice. The S1P1R agonist FTY720 (Sigma-Aldrich) was dissolved in DMSO at a concentration of 20 mg/ml; 1 mg/kg was administered intraperitoneally daily, starting days before influenza A/X31 infection until the mice were sacrificed after viral rechallenge. Mice were anesthetized intraperitoneally with ketamine/xylazine (Patterson Veterinary) and retro-orbital intravenous injection of 3 μg of CD45.2 (104, BioLegend) was then performed to enable flow cytometric differentiation of circulating immune cells in lung vasculature from tissue-resident immune cells in lung parenchyma as previously described.^63^

### Tissue processing

Harvested lung lobes were minced with scissors and digested at 37°C for 30 minutes in RPMI 1640 with Liberase and DNase (Roche). Lung tissue was then strained through a 70 μm filter and subjected to RBC lysis with Red Blood Cell Lysing Buffer (E-bioscience) to generate a single-cell suspension. Mediastinal lymph nodes were harvested from the same mice as indicated and mechanically dissociated through a 70 μm filter and subjected to RBC lysis to generate a single cell suspension.

### Flow cytometry

For surface staining, single cell suspensions were subjected to Fc-receptor blockade with CD16/CD32-blocking antibody (clone 93) (Invitrogen) and were incubated for 30 minutes at 4°C with the indicated anti-mouse antibody. After staining, dead cells were excluded by Fixable Viability Dye eFluor 780 (ThermoFisher). MHC class II tetramer staining was performed by incubating cells for 1 h at 37°C with the NP_311–325_ peptide-loaded IA^b^ tetramer (NIH tetramer core facility) in RPMI 1640 supplemented with 10% FCS. Cells were then washed and subjected to surface staining and viability dye. To label NP-specific B cells, recombinant influenza NP (Sino Biological) was conjugated with APC dye according to the manufacturer’s protocol (Abcam) and added to the B cell antibody solution.

For transcription factor staining, cells were stained for surface markers as documented above, washed, and stained with viability dye. Cells were then fixed and permeabilized with the Foxp3/Transcription Factor Staining buffer set according to the manufacturer’s instructions (eBioscience) and stained with the indicated antibodies for 30 minutes at room temperature.

B cells were stained with CD19 FITC (clone D3/CD19), CD38 BUV396 (clone 90/CD38), GL7 PE (clone GL7), CXCR3 BV421 (clone CXCR3-173), IgD PE-Cy7 (clone 11-26c), and IgM PerCP-Cy5.5 (clone 11/41). T cells were stained with CD4 BV605 (clone 6121.5), CD3 BUV396 (clone 17A2), CD44 PE (clone IM7), CD69 APC (clone H1.2.F3), and FR4 PerCP-Cy5.5. FoxP3 (clone FJ12-165).

Flow cytometry data was acquired using Fortessa X-20 (BD Biosciences) or LSR II (BD Biosciences) instruments with FACSDiva v 8.0.3 software. Data were analyzed using FlowJo v 10.10 software (Waters Corporation).

### Immunofluorescence staining and microscopy

Dissected lung lobes were fixed in 4% paraformaldehyde in PBS overnight at 4 °C and dehydrated in 30% sucrose solution for 48 h before embedding in O.C.T. (Sakura). 12–50 μm cryosections of tissues were collected on Superfrost Plus^TM^ microscope slides, blocked and permeabilized with 5% normal donkey serum (Jackson ImmunoResearch, 017-000-121) made with 0.3% Triton X-100 in PBS for 30 minutes, incubated with primary antibodies overnight at 4 °C and then secondary antibodies for 45 minutes at room temperature after washing with PBS.

Microscopic images were obtained with an Akoya Biosciences Vectra slide-scanning microscope or with a Leica SP8 confocal laser-scanning microscope using a 60x oil, 40x water or 10x air objective. Image processing and analysis were done using Fiji software (Version 1.0) and QuPath 0.7.0-x64 (Version 1).

### Viral quantification

Viral titers were determined by quantification of viral transcripts using reverse transcription–quantitative PCR (RT-qPCR) as previously described.^68,69^ RNA was isolated from right upper lung lobe tissue homogenates prepared in TRIzol (Invitrogen). RNA concentration and purity were assessed using a NanoDrop spectrophotometer (Thermo Scientific, Nanodrop 2000). 1 μg of RNA was reverse transcribed to make cDNA using MultiScribe Reverse Transcriptase (Applied Biosystems). qPCR was performed using FastStart SYBR Green Master Mix (Sigma-Aldrich) on a CFX96 Real-Time PCR Detection System (Bio-Rad) and normalized to GAPDH. Relative mRNA levels were calculated using normalization based on housekeeping genes using the ΔΔCt method.^70^

### Primers

GAPDH, 5’-AGGTCGGTGTGAACGGATTTG-3’ (forward) and 5’-TGTAGACCATGTAGTTGAGGTCA-3’ (reverse) primers and Non-structural protein-1 (NS1), 5’-TGTCAAGCTTTCAGGTAGATTG-3’ (forward) and 5’-CTCTTAGGGATTTCTGATCTC-3’ (reverse) primers.

### Enzyme-linked immunosorbent spot (ELISpot) assay

Multiscreen IP HTS filter plates were coated with 2 μg/ml of influenza nucleoprotein (NP) in carbonate buffer and incubated overnight at 4 °C. Plates were washed with wash buffer (0.05% Tween in PBS) and blocked with 1% BSA in PBS for 1h at 37 °C. Serial dilutions of single-cell suspensions prepared in complete medium (RPMI 1640, 10% fetal bovine serum, 2% L-Glutamine (200 mM), 1% Penicillin-Streptomycin (100X), 1% Sodium Pyruvate (100 mM), 0.1% Beta-mercaptoethanol (50 mM)) were plated and incubated at 37°C in 5% CO_2_ for 16 h. Plates were washed and incubated with goat anti-mouse alkaline phosphatase-conjugated immunoglobulin A (IgA) or goat anti-mouse alkaline phosphatase-conjugated kappa antibody (IgG) diluted (1:1,000) in 0.5% BSA, 0.05% Tween in PBS for 1 h at 37 °C in 5% CO_2_. Plates were washed and developed with BCIP/NBT substrate for 1 h. Spots were recorded using a CTL Immunospot S6 Macroplate Imager Reader and counted manually.

### Profiling influenza-infected lung samples using Slide-tags

Mouse lungs at 2, 3 and 4 weeks post-influenza infection were used for Slide-tags. Fresh-frozen tissues were cryo-sectioned at 20 μm thickness using a cryostat set at −17 to −18 °C. Pre-cooled, custom-made 6-mm square biopsy punches were used to isolate regions of interest from tissue sections. These regions were selected based on checking tdTomato-positive fluorescence signals in an adjacent slide to enrich for M cells. The isolated tissue regions were carefully transferred onto a precooled array, which was placed on top of a glass slide. A finger was briefly placed underneath the slide to melt the tissue onto the array. Immediately after, the tissue, array, and slide were placed on ice, and approximately 10 μl of dissociation buffer (82 mM Na_2_SO_4_, 30 mM K_2_SO_4_, 10 mM glucose, 10 mM HEPES, 5 mM MgCl_2_) was gently pipetted onto the tissue to ensure it was fully covered. The array was then exposed to an ultraviolet (UV) light source (0.42 mW mm^-2^, Thorlabs, M365LP1-C5, Thorlabs, LEDD1B) for 1 minute to cleave spatial barcode oligonucleotides off the beads. After photo-cleavage, the array was incubated on ice for 7.5 minutes before being transferred to a well of a 12-well plate. To release the tissue from the array, 1 ml of extraction buffer (dissociation buffer with 1% Kollidon VA64, 0.2% Triton X-100, 1% BSA, 666 U ml^-1^ RNase inhibitor) was gently dispensed onto the array, and the buffer was carefully triturated up and down over the tissue. This process was repeated until the tissue was completely released from the array. The array was then discarded, and mechanical dissociation of the tissue was performed by triturating the supernatant 100–150 times using a 1 ml pipette to fully release the nuclei from the tissue. The extraction buffer containing the nuclei was transferred to a 15 ml tube. The well was washed three times with 1 ml of wash buffer (dissociation buffer with 1% BSA and 1: 100 RNase inhibitor) and the washes were pooled into the same 15 ml tube. The final volume of the wash buffer was adjusted to 10 ml. The nuclei were centrifuged at 600g for 10 minutes at 4 °C. After centrifugation, 9.5 ml of the supernatant was carefully removed. The pellet was resuspended and passed through a precooled 40 μm cell strainer (Corning, 431750) into a 1.5ml eppendorf tube. The 15 ml tube and cell strainer were washed with 1 ml of wash buffer, and the nuclei were pelleted again by centrifuging at 600g for 10 minutes at 4 °C. After centrifugation, the supernatant was carefully removed, leaving approximately 50 μl of wash buffer for nuclei resuspension. To determine the cell count, 2 μl of resuspended nuclei was mixed with 18 μl of 1:100 diluted DAPI, and the nuclei were manually counted using a C-Chip Fuchs-Rosenthal disposable hemocytometer (INCYTO, DHC-F01-5). Based on the cell count, up to 25,000 nuclei were processed using the Chromium Next GEM Single Cell 5’ Kit v2 (10x Genomics, PN-1000263) with Feature Barcode technology for Cell Surface Protein and Immune Receptor Mapping, following the manufacturer’s protocol with one modification. During cDNA amplification (Step 2.2a), cDNA Primer (10x Genomics, PN-2000089) was used in place of Feature cDNA Primers 4 (PN-2000277), with the addition of 1 μL of a custom primer (5′-GTGACTGGAGTTCAGACGT-3′, 0.3 μM). BCR libraries were amplified from cDNA and prepared for Illumina sequencing according to the 10x Genomics workflow.

## QUANTIFICATION AND STATISTICAL ANALYSIS

### Statistical Analysis

Statistical analyses were performed using GraphPad Prism 11 (GraphPad Software, Inc.). The numbers of mice and replicates are indicated in the corresponding figure legends. Statistical significance was assessed using unpaired, two-tailed Student’s *t*-tests, Welch’s *t*-tests, Mann–Whitney U tests, or ordinary one-way analysis of variance (ANOVA), as appropriate. Tukey’s multiple-comparisons test was applied following one-way ANOVA. The statistical tests used for individual experiments are specified in the figure legends. Statistical significance was defined as *p* < 0.05, with significance levels indicated as follows: *p* < 0.05 (*), *p* < 0.01 (**), *p* < 0.001 (***), and *p* < 0.0001 (****).

### Slide-tags data preprocessing

BCL files from transcriptome libraries were demultiplexed into FASTQ files using the cellranger mkfastq command with default parameters (Cell Ranger v7.0.0; 10x Genomics). Transcriptome reads were aligned to the mouse mm10 reference genome (GENCODE vM23/Ensembl 98) using cellranger count, with intronic reads included using the –include-introns option. Gene expression counts were quantified as unique molecular identifiers (UMIs). Spatial barcode libraries were demultiplexed into FASTQ files using *bcl2fastq* and were then processed through the Slidetags pipeline (https://github.com/broadchenf/Slide-tags) to calculate spatial locations for nuclei.

### Cell-type annotation

Data from different libraries derived from the same tissue section were processed independently using Cell Ranger and the Slide-tags pipeline described above. Cells were retained if spatial location information was available and were filtered to exclude those with fewer than 200 detected genes, gene or UMI counts above the 99.9th percentile, or greater than 10% mitochondrial transcript content. Filtered Seurat (v5.0.0) objects from different tissue sections were then merged. The merged dataset was log-normalized, and the top 2,000 variable features were identified using FindVariableFeatures, followed by principal component analysis using RunPCA. The first 50 principal components were used for UMAP visualization and graph-based clustering using RunUMAP, FindNeighbors, and FindClusters (dims = 1:50; resolution = 0.8). Broad cell populations, including epithelial, endothelial, fibroblast, immune, and mesothelial cells, were initially annotated based on established lineage-specific marker genes. Each major population was then iteratively subclustered to enable more refined annotation of distinct cell types and states.

### Spatial neighborhood enrichment analysis

Cell-type co-localization was assessed in Slide-tags data using cell-type label permutation. For each tissue section, a fixed-radius adjacency graph was constructed using frNN in dbscan (v1.1.12), excluding self-neighbors. Cells with no neighbors within the search radius were excluded from enrichment calculations. For each pair of cell types, the fraction of type-j neighbors surrounding type-i center cells was calculated using sparse matrix operations implemented with *Matrix* (v1.6.5). To generate an empirical null distribution, cell-type labels were independently permuted 1,000 times within each tissue section, thereby preserving the spatial coordinates and cell-type composition of each section, and neighbor fractions were recalculated for each permutation. For sections of the same harvest time point, the observed neighbor fractions and those from each permutation were aggregated across sections as weighted means, with weights proportional to the number of center cells of each type per section. For each cell-type pair, an enrichment Z-score was then calculated as the difference between the observed enrichment and the mean of the corresponding permutation-derived null distribution, normalized by its standard deviation. Median neighbor counts per cell-type pair were computed using matrixStats (v1.1.0).

To determine the analysis radius, the permutation analysis was repeated across a range of neighborhood radii (50–500 µm), and pairwise Spearman correlations were calculated between vectors of cell-type-pair enrichment Z-scores obtained at each radius. Enrichment patterns were highly consistent across the tested radii. A radius of 200 µm was selected for the primary analysis based on a balance between enrichment signal strength and spatial interpretability.

### BCR processing

V(D)J alignment and clonotype assembly were performed with MiXCR (v4.7.0) using the 10x-sc-xcr-vdj preset for mouse (Mus musculus). Heavy-chain (IGH) clonotypes were exported with top V, D, and J segment calls, CDR3 nucleotide and amino acid sequences (Table S1).

Downstream processing was restricted to cells present in quality-controlled B and plasma cells (including cells with and without spatial information). Only heavy-chain clonotypes were retained (IgM, IgG, IgA, IgE), and CDR3 sequences shorter than 9 nucleotides (3 amino acids) were excluded. For cells with multiple heavy-chain reads, a single chain was selected by matching each candidate isotype to the corresponding Ig constant-region gene expression in the scRNA-seq data and retaining the isotype with the highest expression; in cases where expression was uninformative, the chain with the highest UMI count was kept. The primary analysis used a UMI ≥ 1 threshold; a UMI ≥ 2 filter was applied as a sensitivity check.

Clonal families were assigned in two stages. First, cells were grouped by shared V, D, and J segments. Within each V/D/J group, cells were further stratified by CDR3 nucleotide length. Within each V/D/J and length stratum, pairwise Levenshtein distances between CDR3 nucleotide sequences were computed and normalized by CDR3 length to yield a fractional mismatch. Single-linkage hierarchical clustering with a threshold of 0.20 (≤20% sequence divergence) was applied to define clonal families, implemented using stringdist (v1.9).

Clonal families were further classified by intra-family CDR3 diversity. Families containing two or more cells with identical CDR3 amino acid sequences and no additional variants were classified as clones, representing a single progenitor B cell that proliferated without further CDR3 diversification. Families containing two or more distinct CDR3 amino acid sequences were designated clonal families, indicating diversification through somatic hypermutation. Cells not grouped with any other cell were classified as singletons (Table S1).

### Intra-family CDR3 sequence divergence and isotype concordance

For each family, all pairwise combinations of plasma cells were enumerated. For each pair, CDR3 sequence divergence was quantified as the normalized Levenshtein distance between nucleotide CDR3 sequences, and isotype concordance was defined as a binary indicator of whether the two cells shared the same broad isotype class (IgA or IgG). Cell pairs were binned by CDR3 divergence into 0–0.05, 0.05–0.10, 0.10–0.15, 0.15–0.20 and mean isotype concordance was calculated within each bin. Two reference baselines were defined. The empirical random null (0.50) represents the expected concordance if isotypes were assigned independently at random according to the observed isotype frequency distribution. Complete isotype conservation (1.0) represents the expectation where all plasma cells carry the same isotype regardless of CDR3 divergence. Statistical significance at each divergence bin was assessed by one-sided binomial test against the empirical null.

### Spatially resolved ligand-receptor analysis through LARIS

Spatially resolved ligand-receptor interactions were identified using LARIS (v0.12.0) in cell-type-specific mode and k = 20 spatial nearest neighbors. For visualization in Figure 6A, cell-type-specific interaction scores were filtered to interactions involving M cells as senders and immune cell types as receivers with FDR < 0.05 (Table S2).

### Spatial visualization of M-immune cell interaction

For visualization of specific M-immune cell interactions at single-cell resolution, a spatial kNN (k = 20) was constructed from cell coordinates. For visualization of *RANKL*-*RANK* signaling to M cells, Tfh, Treg, and Th17 cells were identified as candidate senders if at least one M cell was present among their k = 20 nearest spatial neighbors. Candidate sender cells are displayed with dot size proportional to their own *Tnfsf11* (*RANKL*) expression. M cells are displayed with dot size proportional to their own *Tnfrsf11a* (*RANK*) expression. For visualization of M cell secreting cytokines, sender scores were computed for each M cell as the product of its ligand expression (*Ccl20*, *Flt3l* and *Il7*) and the mean receptor expression (*Ccr6*, *Flt3* and *Il7r* respectively) across its k nearest immune-cell neighbors, capturing local paracrine signaling capacity. Receiver scores were computed for each immune cell as the product of its receptor expression and the mean ligand expression across its k nearest M-cell neighbors, reflecting proximity-weighted exposure to M cell-derived ligand. Scores were mapped onto tissue coordinates, with dot size proportional to score magnitude, allowing direct visualization of spatially active sender and receiver cells within and around iBALT structures.

