## Supplementary figures and images for "Induced microfold cells are required for the function of pulmonary tertiary lymphoid organs that promote mucosal immunity"

### Supplemental Figure 1

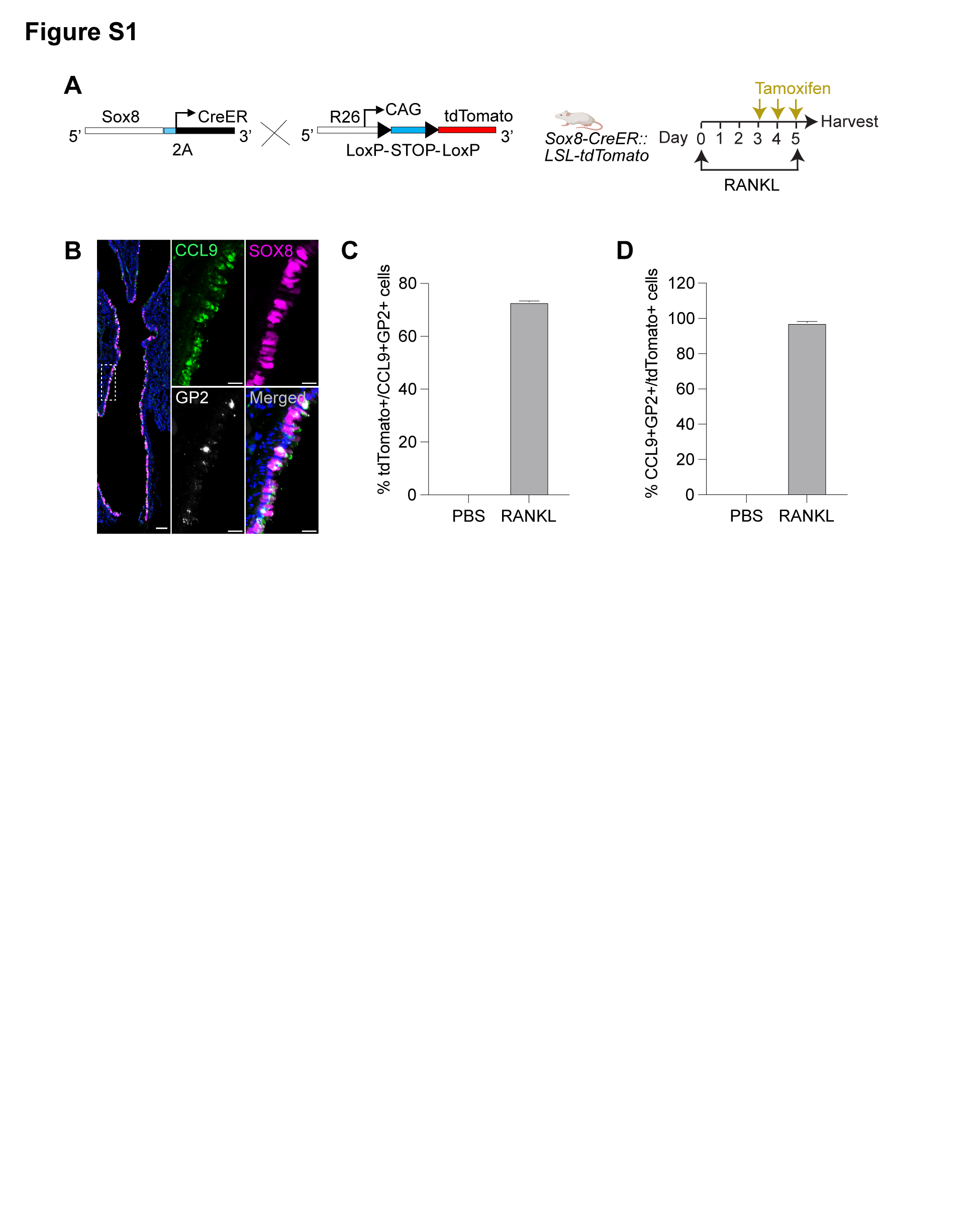

### Supplemental Figure 2

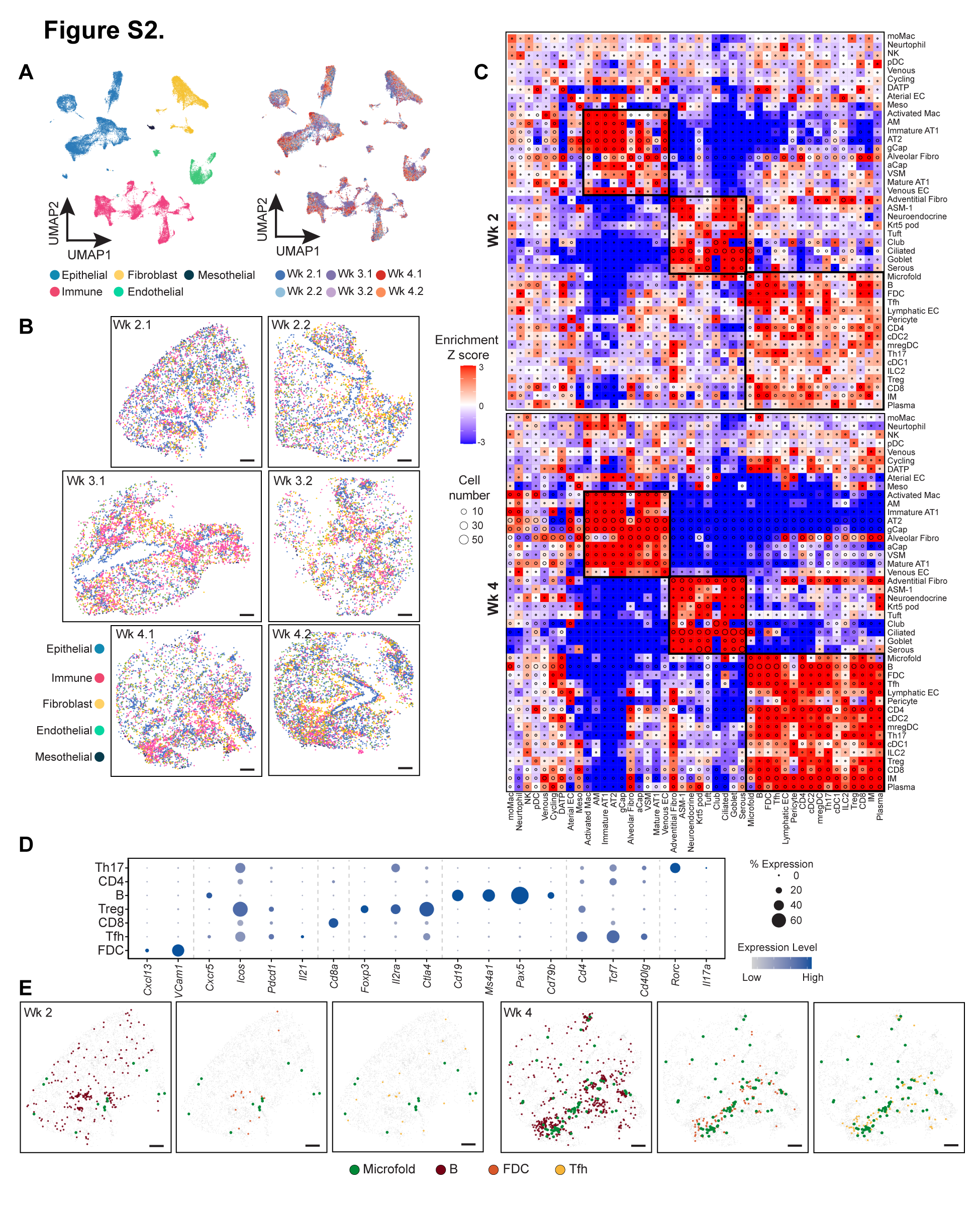

### Supplemental Figure 3

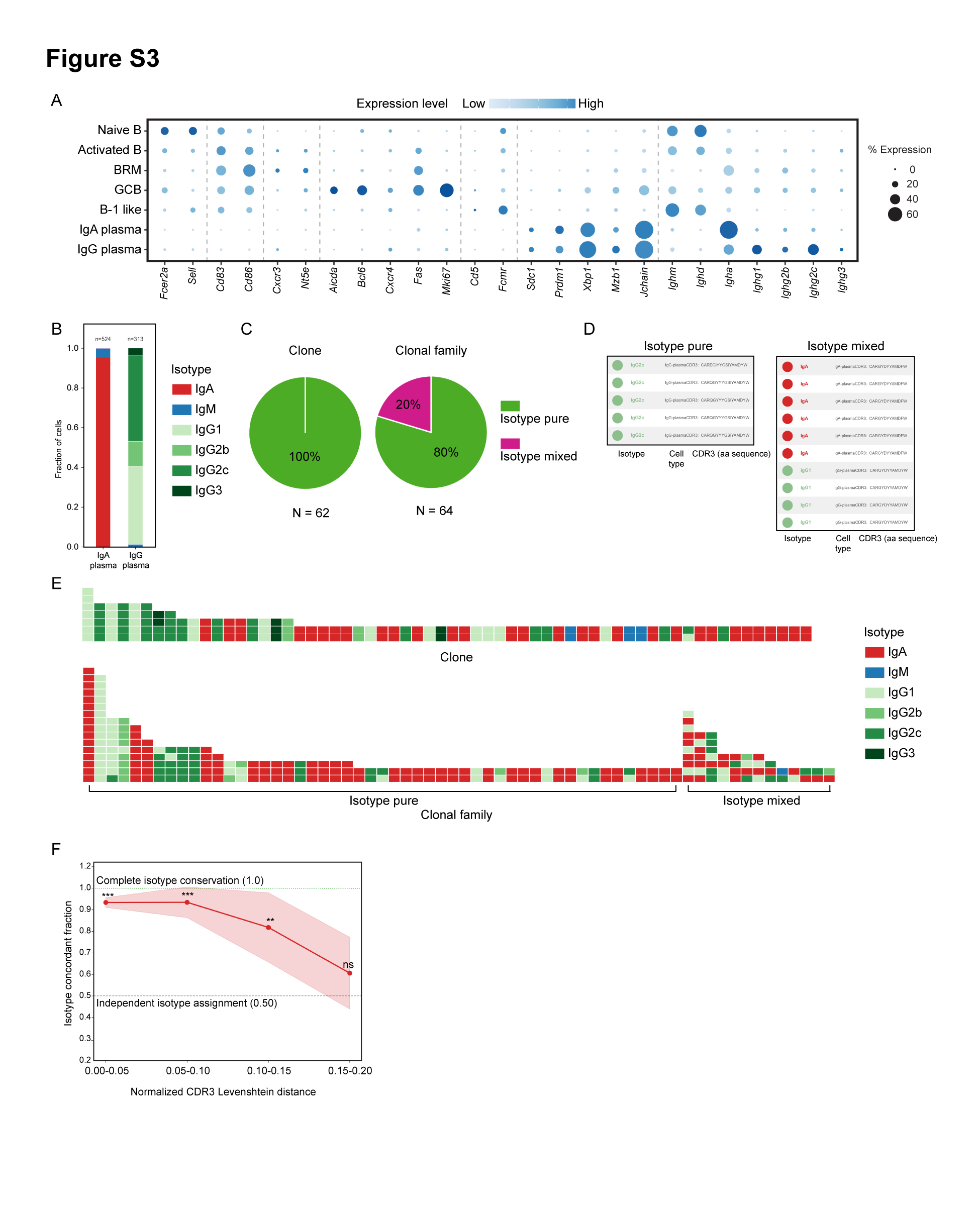

### Supplemental Figure 4

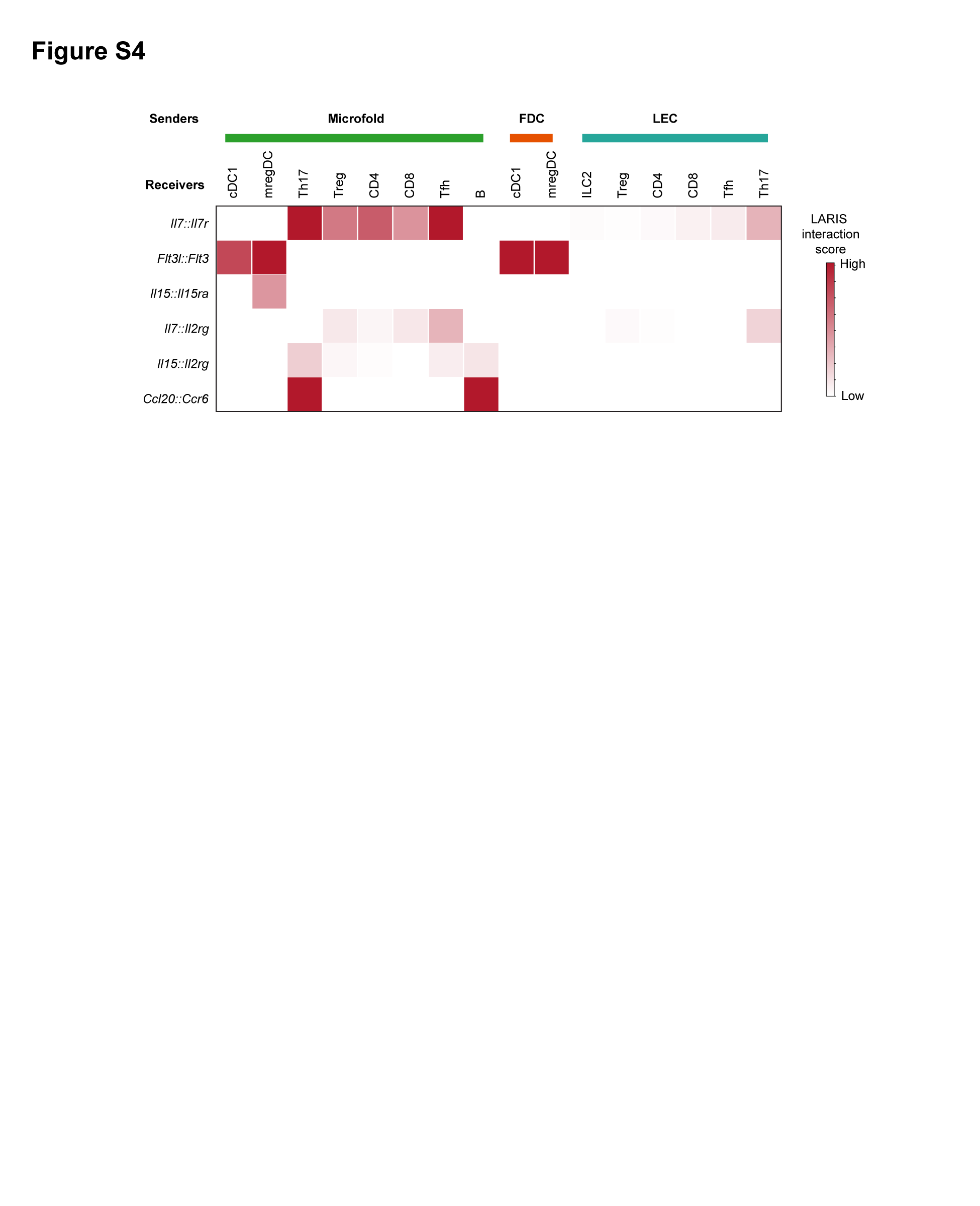

### Supplemental Figure 5

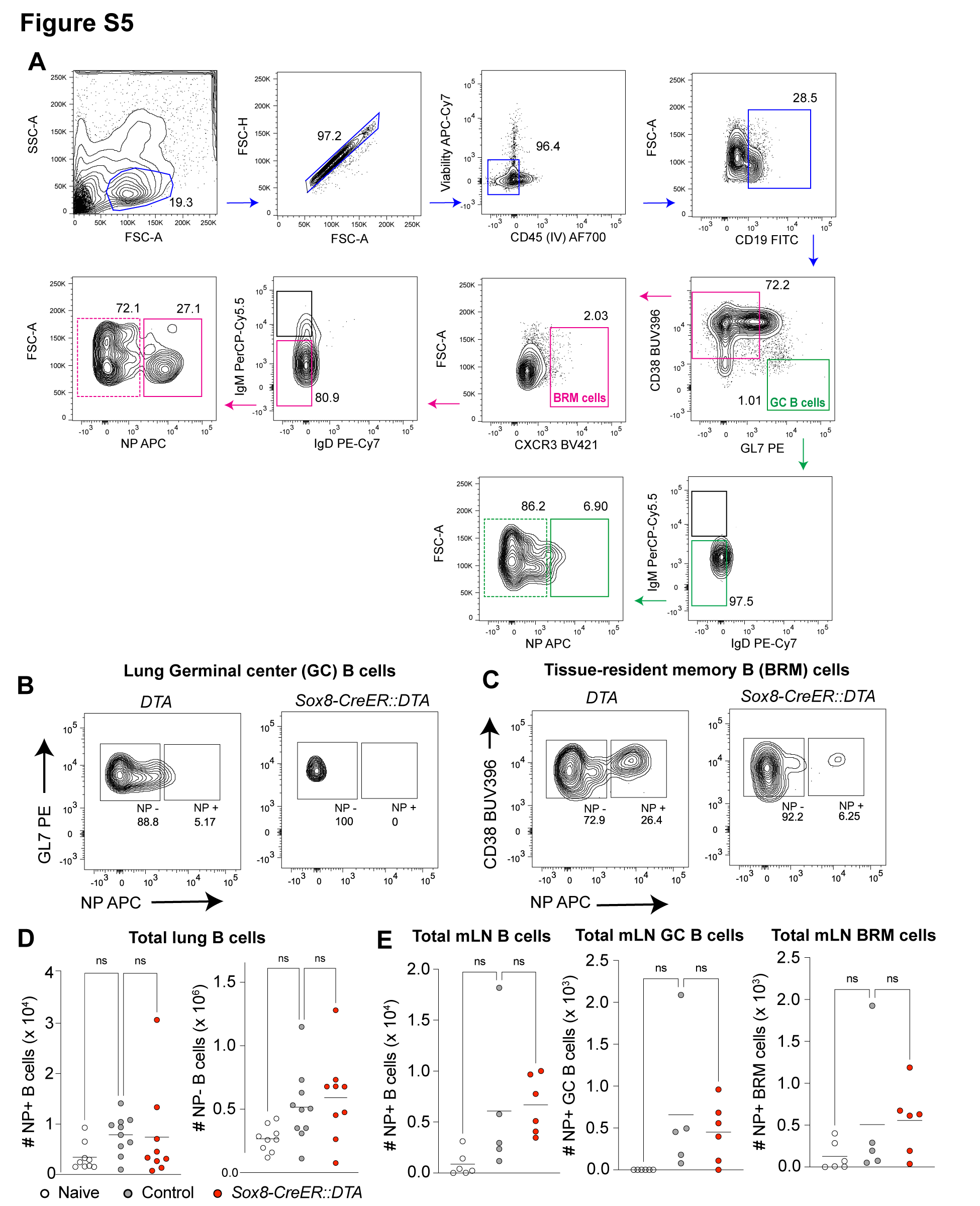

### Supplemental Figure 6

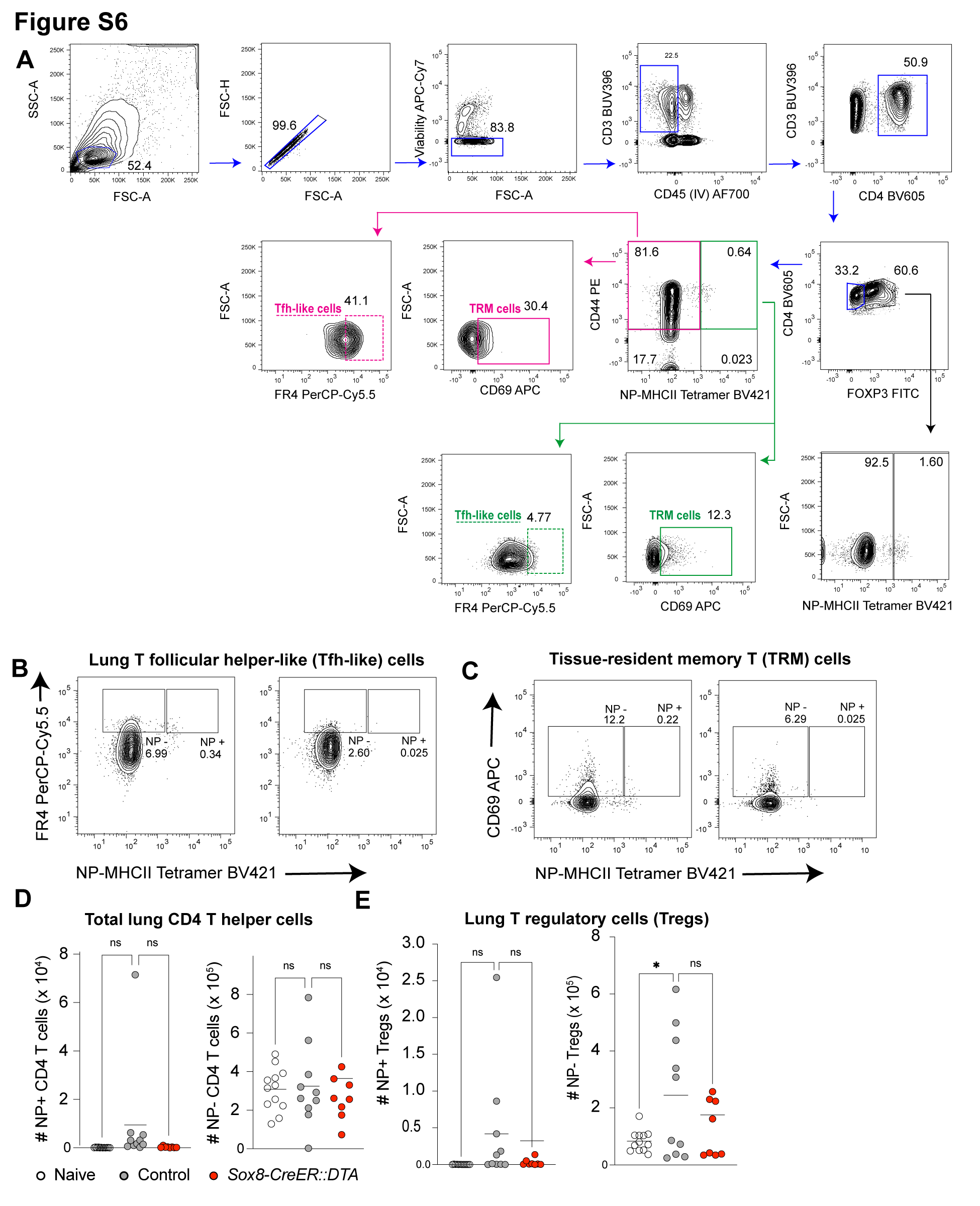
